# Benchmarking deep learning methods for biologically conserved single-cell integration

**DOI:** 10.1101/2024.12.09.627450

**Authors:** Chenxin Yi, Jinyu Cheng, Wanquan Liu, Junwei Liu, Yixue Li

## Abstract

Advancements in single-cell RNA sequencing (scRNA-seq) have enabled the analysis of millions of cells, but integrating such data across samples and methods while mitigating batch effects remains challenging. Deep learning approaches address this by learning biologically conserved gene expression representations, yet systematic benchmarking of loss functions and integration performance is lacking. This study evaluated 16 integration methods using a unified variational autoencoder framework, incorporating batch and cell-type information. Results revealed limitations in the single-cell integration benchmarking index (scIB) for preserving intra-cell-type information. To address this, we introduced a correlation-based loss function and enhanced benchmarking metrics to better capture biological conservation. Using annotations from the Human Lung Cell Atlas and Human Fetal Lung Cell Atlas, our approach improved biological signal preservation. This work highlights the need for biologically informed metrics in scRNA-seq integration and offers guidance for future deep learning developments.

## Main

Single-cell RNA sequencing (scRNA-seq) has revolutionized our ability to study cellular diversity by providing high-resolution insights into gene expression at the single-cell level^1^. With the advancement of scRNA-seq techniques, the volume of single-cell data across various species, tissues, and developmental stages has expanded significantly, growing from hundreds of cells to tens of millions^2^. The scRNA-seq data analysis aims to understand cellular gene expression and functional variations across different diseases or developmental stages, helping to identify mechanisms underlying cell alterations^3^. Among that, data integration is a critical process for combining data collected from different samples and time points and is also essential for incorporating external data with a similar biological background^4^. However, due to potential data biases, high data dimension, and sparsity in scRNA-seq data, integrating large-scale single-cell data from different experiments, studies, and platforms while preserving crucial biological insights remains a significant challenge^5, 6^.

Several statistical methods have been developed to address batch effects in scRNA-seq data integration. One strategy is to identify the Mutual Nearest Neighbors (MNN) of single cells across datasets, including MNN^7^, Scanorama^8^, and Seurat V3^9^. Another strategy focuses on balancing cellular neighbors to prevent batch-specific clustering, exemplified by Harmony^10^ and batch-balanced k-nearest neighbors (BBKNN)^11^. Additionally, Non-Negative Matrix Factorization (NMF) is employed to identify datasets-shared factors for integration, as seen in LIGER^12^. scMerge^13^ and scMerge2^14^ leverage stably expressed genes or pseudo-replicates to estimate the factor of interest and mitigate unwanted batch variation. While these methods are effective in aligning and integrating scRNA-seq data, they often encounter difficulties with large-scale datasets, particularly those exhibiting high cell-type heterogeneity across datasets^15^.

Deep learning-based approaches have emerged as more powerful and flexible solutions for single-cell data integration^16^. The capability of deep learning methods in learning large, high-dimensional, and complex datasets advances its ability to obtain crucial biological variation^17^. Autoencoder is a versatile framework to learn the latent data representation of high-dimensional single-cell gene expression data. Li et al. developed the DESC method, which employs an autoencoder to infer unsupervised embedding of scRNA-seq data and for batch-invariant cell clustering analysis^18^. Another notable method is single-cell Variational Inference (scVI)^19^, a fully probabilistic deep learning framework that accounts for both biological and technical noise in scRNA-seq data. scVI uses a conditional variational autoencoder (cVAE) framework to treat different batches as variables while preserving true biological gene expression information. Moreover, the deep learning framework facilitates more complex model designs and enhances information regularization. The SCALEX method introduces a batch-free encoder to project the batch-invariant embeddings across datasets^20^. Additionally, in atlas-level data integration, pre-defined cell type information can also be leveraged for data integration. Single-cell ANnotation using Variational Inference (scANVI)^21^ extends scVI by incorporating pre-existing cell state annotations, improving the accuracy of cell type identification in new datasets through a semi-supervised learning approach. Similarly, methods including scDREAMER^22^ and scDML^23^ can also utilize predefined cell clustering information for semi-supervised batch-removal data integration.

The success of deep learning methods in single-cell data batch correction and integration largely depends on the design of the loss function. Generally, the objective of these methods is to remove unwanted batch effects while preserving biological information across single-cell datasets. Both batch effects and biological signals can be partially captured by batch labels and predefined cell-type labels, respectively. To address batch correction, techniques such as adversarial learning^22, 24^ and information-constraining^25^ methods are employed to minimize batch-specific information across datasets. For preserving biological information, strategies like supervised domain adaptation^26^ and deep metric learning^23^ are utilized to ensure cell-type label information is maintained in the integrated data. While various loss function designs are effective in different methods, a horizontal comparison of the impact of distinct loss function combinations in single-cell data integration tasks is lacking. Additionally, in the context of single-cell integration performance benchmarking, the single-cell integration benchmarking (scIB)^27^ framework primarily evaluates methods in two key areas: batch correction and biological conservation, based on the batch and cell-type labels. While scIB provides a robust foundation for performance evaluation, it falls short in fully capturing unsupervised intra-cell-type variation. As deep learning models continue to evolve, there is a growing need for more refined benchmarking metrics that can accurately assess both batch effect correction and the preservation of critical biological information.

In this study, we developed 16 deep-learning single-cell integration methods across three distinct levels within a unified variational autoencoder framework. These methods were designed to comprehensively evaluate the impact of different loss function combinations on data integration, utilizing batch information, cell-type information, or both jointly. By analyzing the effects of batch correction and biological conservation across varying loss function configurations, we identified that current benchmarking metrics and batch-correction methods fail to adequately capture intra-cell-type biological conservation. This finding was validated with multi-layered annotations from the Human Lung Cell Atlas (HLCA)^28^ and the Human Fetal Lung Cell Atlas^29^. To address this gap, we introduced a correlation-based loss function to better preserve biological signals and refined existing benchmarking metrics by incorporating intra-cell-type biological conservation. Our findings highlight the potential of deep learning methods for single-cell data integration, with the refined framework and benchmarking metrics offering deeper insights into the integration process. These advancements are poised to drive the development of deep learning methods for integrating increasingly complex multimodal and spatiotemporal single-cell data.

## Results

### Deep-learning-based integration of single-cell data

Single-cell data integration is essential for atlas-level single-cell data analysis, and the advent of deep learning methods has broadened the application of data integration, enabling a deeper understanding of diverse biological processes. In this study, we present a unified benchmarking framework that evaluates different loss function designs and information regularization strategies for the data integration task (Figs. 1a, b). We also reorganize existing benchmarking metrics and expand their applications to include batch correction and biological conservation at both inter-cell-type or intra-cell-type levels (Fig. 1c). Additionally, we validate our extended single-cell integration benchmarking (scIB-E) metrics with multi-layer cell annotation and developmental single-cell atlas datasets, and incorporated a novel loss function designed for intra-cell-type biological conservation (Fig. 1d).

**Fig 1.**
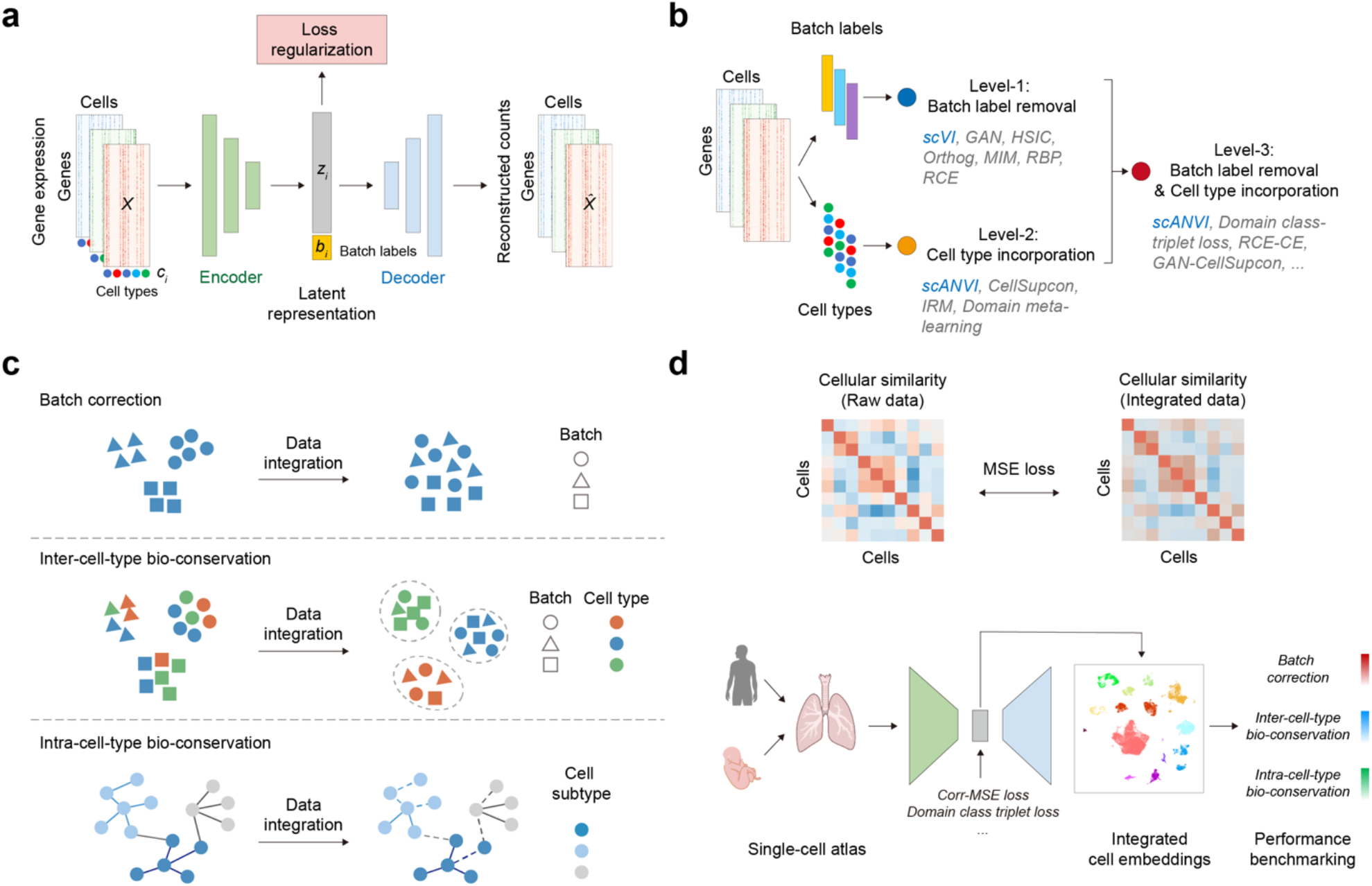
Multi-level loss regularization designs for single-cell integration. **a**, Diagram illustrating the unified variational autoencoder framework used in this project. **b**, Schematic representation of the three-level loss designs implemented in this study. **c**, Illustration of the effects of batch correction (top), inter-cell-type biologic conservation (middle), and intra-cell-type biologic conservation (bottom) following single-cell data integration. **d**, Schematic representation of the Corr-MSE loss design (top) and the process of biologically conserved single-cell integration (bottom).

In this study, we used the scVI^19^ and scANVI^21^ models as the foundational deep-learning framework to evaluate the effectiveness of different loss function designs and regularization modules for single-cell data integration. Both methods utilize conditional variational autoencoders to embed single-cell gene expression data, with batch labels of single-cell samples as conditional variables to remove batch effects. In scANVI, known cell-type labels are incorporated for semi-supervised data integration, enabling the preservation of biological information. Consequently, we applied both batch labels of single-cell samples and known cell-type labels of single cells as the proxies of both batch effects and biological information, respectively. Thereby, we developed a multi-level deep-learning strategy for single-cell data integration (Fig. 1b). Level-1 focuses on batch effect removal using batch labels, level-2 incorporates single-cell cell-type labels as biological conservation restraint, and level-3 integrates both batch and cell-type information for data integration. Overall, we designed 16 data integration methods based on the scVI framework across three levels, with the scVI method serving as the baseline for level-1, and the scANVI method as the baseline for level-2 and -3.

### Multi-level loss function designs for single-cell data integration

The level-1 methods are designed to eliminate batch effects by using the batch labels of different single-cell samples. These methods primarily focus on constraining the information shared between the learned latent embeddings of single-cell data and their batch labels. To achieve this, various loss functions are employed, including Generative Adversarial Network (GAN)^30^, Hilbert-Schmidt Independence Criterion (HSIC)^25^, Orthogonal Projection Loss (Orthog), Mutual Information Minimization (MIM), Reverse Backpropagation (RBP)^31^ and Reverse Cross-Entropy (RCE)^32^ (Methods). The level-2 methods incorporate known cell-type labels as proxy biological information to ensure that the latent embeddings from different batches remain biologically aligned. The loss functions at this level include Cell Supervised contrastive learning (CellSupcon)^33^, Invariant Risk Minimization (IRM)^34^ , and Domain meta-learning^35^ (Methods). In level-3 methods, we integrated both batch label and cell-type label to simultaneously achieve batch-effect removal and biological conservation. Leveraging the flexibility of deep learning frameworks, we combine certain loss functions from level-1 and level-2 for level-3, and introduce an additional Domain class triplet loss^36^ (Methods). The model training process follows the scVI framework, and the hyperparameters for the various methods were determined using the automated Ray Tune^37^ framework (Methods). The hyperparameters used for the loss function combinations for each method are listed in Supplementary Tables 1 and 2.

We utilized multiple single-cell RNA-seq datasets to benchmark the performance of various loss designs, including datasets from immune cells^38^, pancreas cells^39^, and the Bone Marrow Mononuclear Cells (BMMC) dataset originating from the NeurIPS 2021 competition^40^ (Supplementary Table 3). For performance evaluation, we used the single-cell integration benchmarking (scIB)^27^ metrics, which provide quantitative evaluations of single-cell integration methods. The Uniform Manifold Approximation and Projection (UMAP)^41^ was utilized to visualize the learned single-cell embeddings of different methods, highlighting the cell distributions across batches and cell types (Fig. 2a, Supplementary Figs. 1-3). We assessed performance by comparing batch correction and biological conservation scores from scIB across multiple datasets (Fig. 2b1, Supplementary Table 4). Compared to the scVI baseline, most level-1 methods demonstrated improved batch correlation index, particularly in RBP regulation (Fig. 2c). For level-2 and level-3 methods, our results indicated that most methods achieved higher biological conservation scores compared to the scANVI baseline. Notably, the level-3 Domain class triplet loss, RBP-CellSupcon, and RCE-CE loss designs outperformed the baseline at both batch correction and biological conservation levels. Additionally, the level-2 IRM loss demonstrated robust biological conservation capability (Fig. 2c).

**Fig 2.**
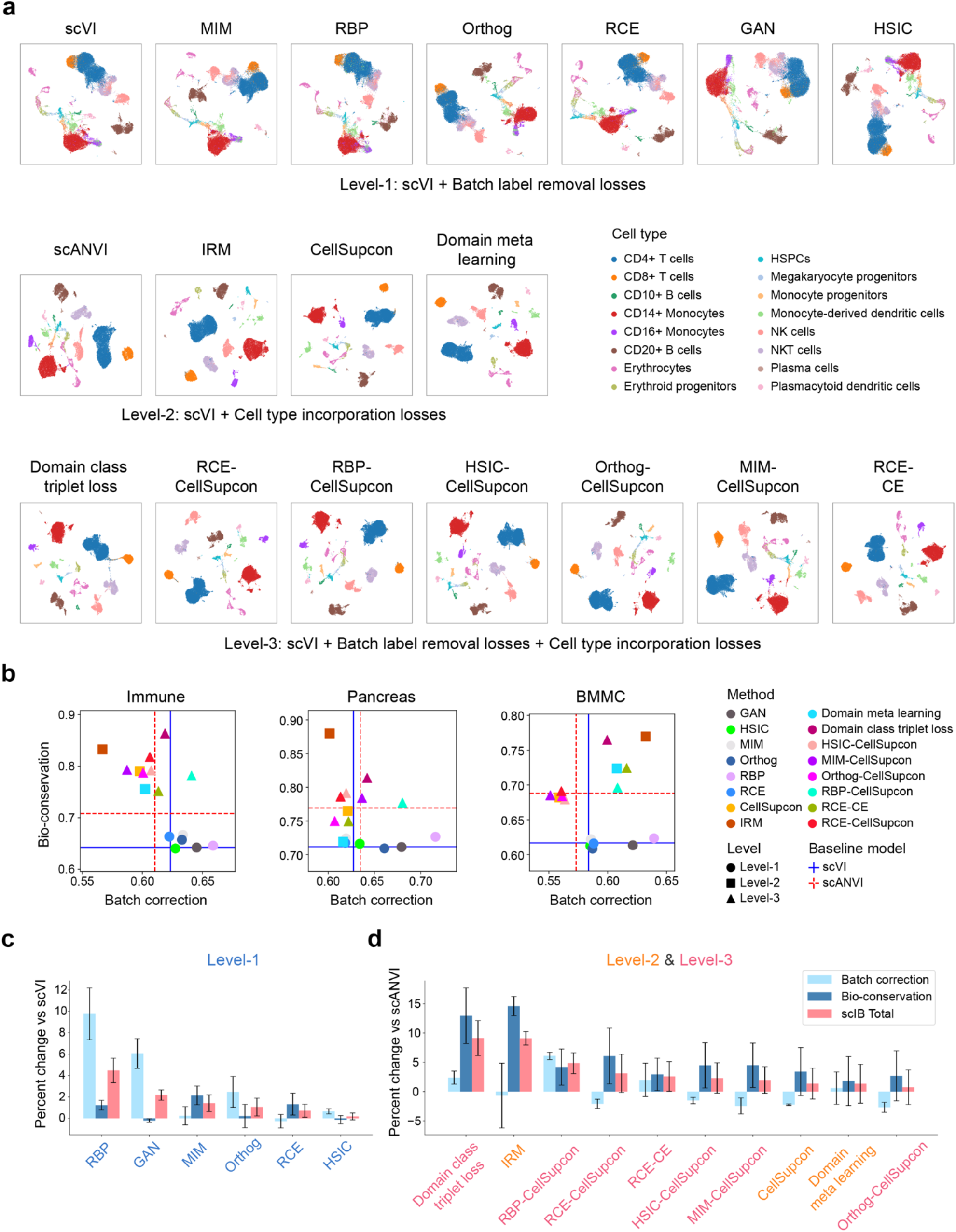
Evaluation of multi-level loss functions for single-cell integration. **a**, UMAP visualization of the immune dataset using three-level integration methods, with cells colored by cell type. **b**, Comparison of the overall batch correction and bio-conservation scores across different methods by three datasets, with point colored by method and shaped by integration levels. Baseline performance of scVI and scANVI are marked by cross lines. **c, d**, Percentage margins of scIB metrics for level-1 (c) and level-2/level-3 (d) methods compared to scVI and scANVI baselines. Bars represent the average percentage margins for different scores, with error bars indicating the standard error of the mean. Methods are ranked in descending order based on their total scIB scores.

### Assessing intra-cell-type biological information in data integration

Based on the performance results across various loss designs and levels of information regularization (Fig. 2), we observed that the strength of information regularization plays a pivotal in single-cell data integration, affecting both scIB metrics for batch correction and biological conservation. To further evaluate the effects of different levels of information regularization, we applied the CellSupcon^33^ loss function with varying hyperparameters (0, 10, 50, and 100), simulating the absence and increasing intensities of cell-type information regularization. Using the same immune dataset^38^, we found stronger cell-type information regularization associated with more distinct separations among annotated single-cell subtypes in the resulting embeddings (Fig. 3a). Moreover, an analysis of scIB metrics across these hyperparameters revealed a gradient increase in the biological conservation score, accompanied by a slight decline in the batch correction score, ultimately resulting in an improvement in the scIB total metric with higher levels of cell-type regularization (Fig. 3b), indicating improved single-cell integration performance.

**Fig 3.**
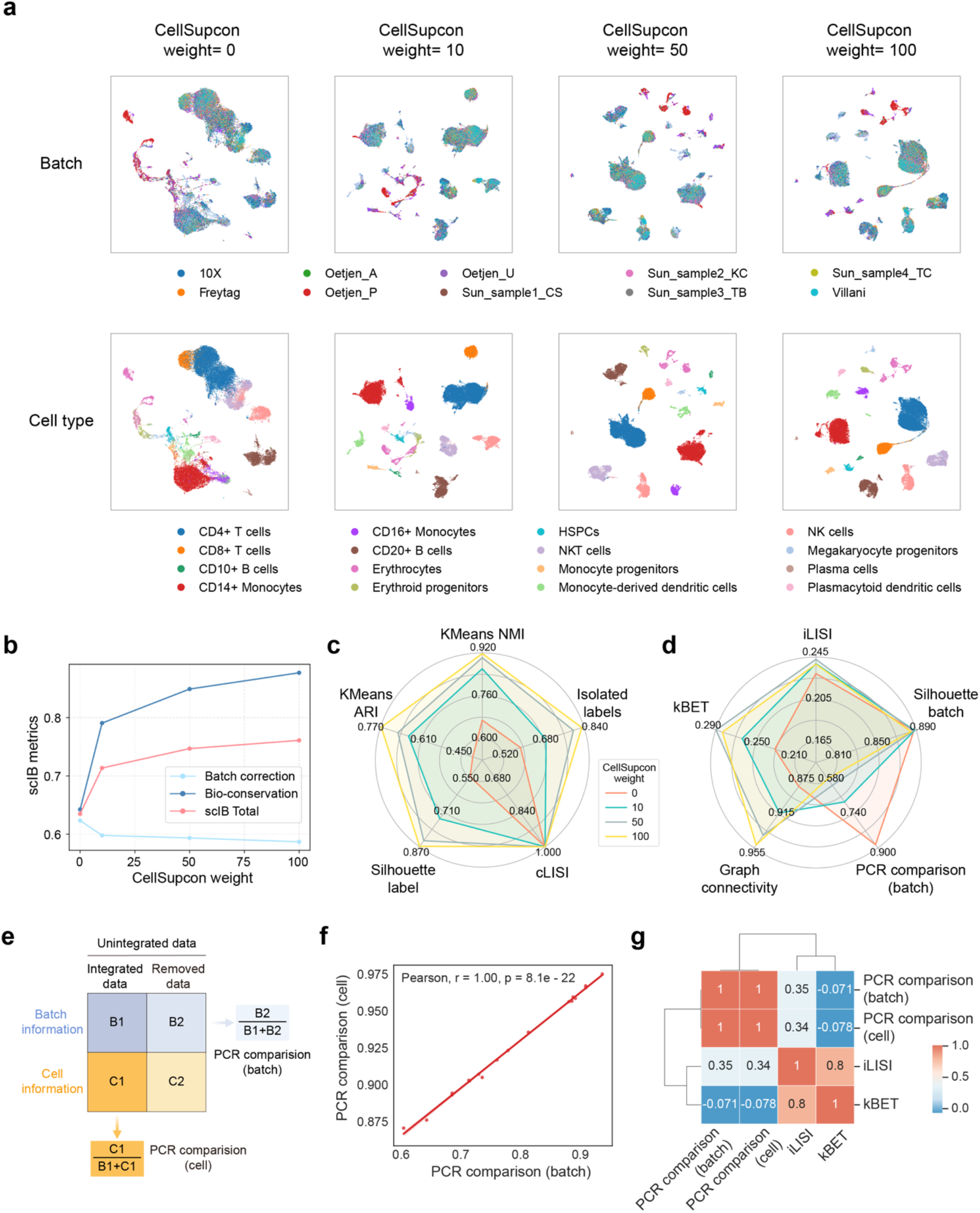
Evaluation of intra-cell-type biological information across varying loss regularizations. **a**, UMAP visualization of the immune dataset using methods with varying weights of the CellSupcon loss, with cells colored by batch label (top) and cell type (bottom). **b**, Line plot showing scIB metrics of single-cell integration results as in (a) with different weights of the CellSupcon loss. **c, d**, Radar plots illustrating trends across detailed indices within biological conservation (c) and batch correction (d) categories of scIB metrics. Colors represent different weights of the CellSupcon loss. **e**, Schematic representation of the design for PCR comparison indices (batch and-cell) to assess biological and batch information across integrated and unintegrated single-cell embeddings. **f**, Pearson correlation analysis between PCR comparison-batch and PCR comparison-cell indices across single-cell integration results, as shown in **Fig. 2a**, for the immune dataset. The p-value is annotated. **g**, Heatmap showing Pearson correlation coefficients between PCR comparison indices (PCR comparison-batch and PCR comparison-cell) and local cellular neighbor-based batch correlation indices (iLISI and kBET).

We then analyzed the trends across specific benchmarking indices within both scIB metric categories. For biological conservation metrics, all indices increased with stronger cell-type information regularization (Fig. 3c, Supplementary Table 5), suggesting that stronger cell-type regularization enhances biological conservation. In the batch correction category, we observed increases in Graph Connectivity and partial improvements in iLISI and kBET scores. However, the Principal Component Regression (PCR) comparison index consistently decreased as cell-type information regularization intensified (Fig. 3d). Graph Connectivity, iLISI, and kBET indices all operate on the k-Nearest Neighbors (kNN) graph constructed from all single-cell data. Specifically, Graph Connectivity quantifies how well cells are connected within annotated cell types, while iLISI and kBET assess the distribution of batch labels across graph nodes (Methods). The observed increases in these indices suggest that embeddings of cells with the same cell-type labels become more similar across batches.

The PCR index quantified the variance in principle components of single-cell embeddings attributable to the batch variable, while the PCR comparison index evaluated the extent of batch information removed after batch correction (Methods). Since the PCR index reflects relative variance in single-cell embeddings, we hypothesized that the PCR comparison index for batch labels (PCR comparison-batch) is potentially inversely related to the preservation of biological information in integrated datasets. To test this, we designed a new PCR comparison index for cellular biological information (PCR comparison-cell), which measures the proportion of cellular variance preserved in integrated single-cell data (Fig. 3e, Methods). Empirical correlation analyses confirmed the consistency between these PCR comparison indices, revealing that a reduction in the PCR comparison-batch with increasing regularization corresponds to a loss of underlying biological information (Fig. 3f). Meanwhile, the PCR comparison indices differed from the other batch correlation indices, which primarily evaluate local cellular neighbors (Fig. 3g, Supplementary Table 6).

### Correlation-based loss function for comprehensive biological conservation

The biological conservation metrics used in scIB are heavily dependent on pre-annotated cell-type labels. However, the PCR comparison-cell index revealed an opposite trend concerning these biological conservation metrics. This led us to hypothesize that the metrics might inadequately capture intra-cell-type biological variation, as they rely solely on the pre-annotated cell-type labels. The intra-cell-type biological variation reflects cellular diversity within annotated populations, encompassing potentially significant biological signals related to micro-populations or disease-specific subtypes.

To confirm that, we utilized a multi-layer annotated single-cell lung atlas featuring five levels of cellular annotations^28^, ranging from broad cell populations to the finest resolved subsets. This framework allowed us to examine single-cell biological information at varying scales. Using the level-1 cell-type labels for model training, we assessed scIB indices across different methods with varying CellSupcon weights. To quantify the conservation of intra-cell-type biological variation, we analyzed the changes in scIB indices across cell annotation levels. Our findings revealed consistent trends in the batch correction index with varying CellSupcon weights across annotation levels (Fig. 4a). However, the biological conservation indices varied across cell levels, suggesting that current biological conservation metrics fail to fully capture cellular biological diversity (Fig. 4a, Supplementary Table 7).

**Fig 4.**
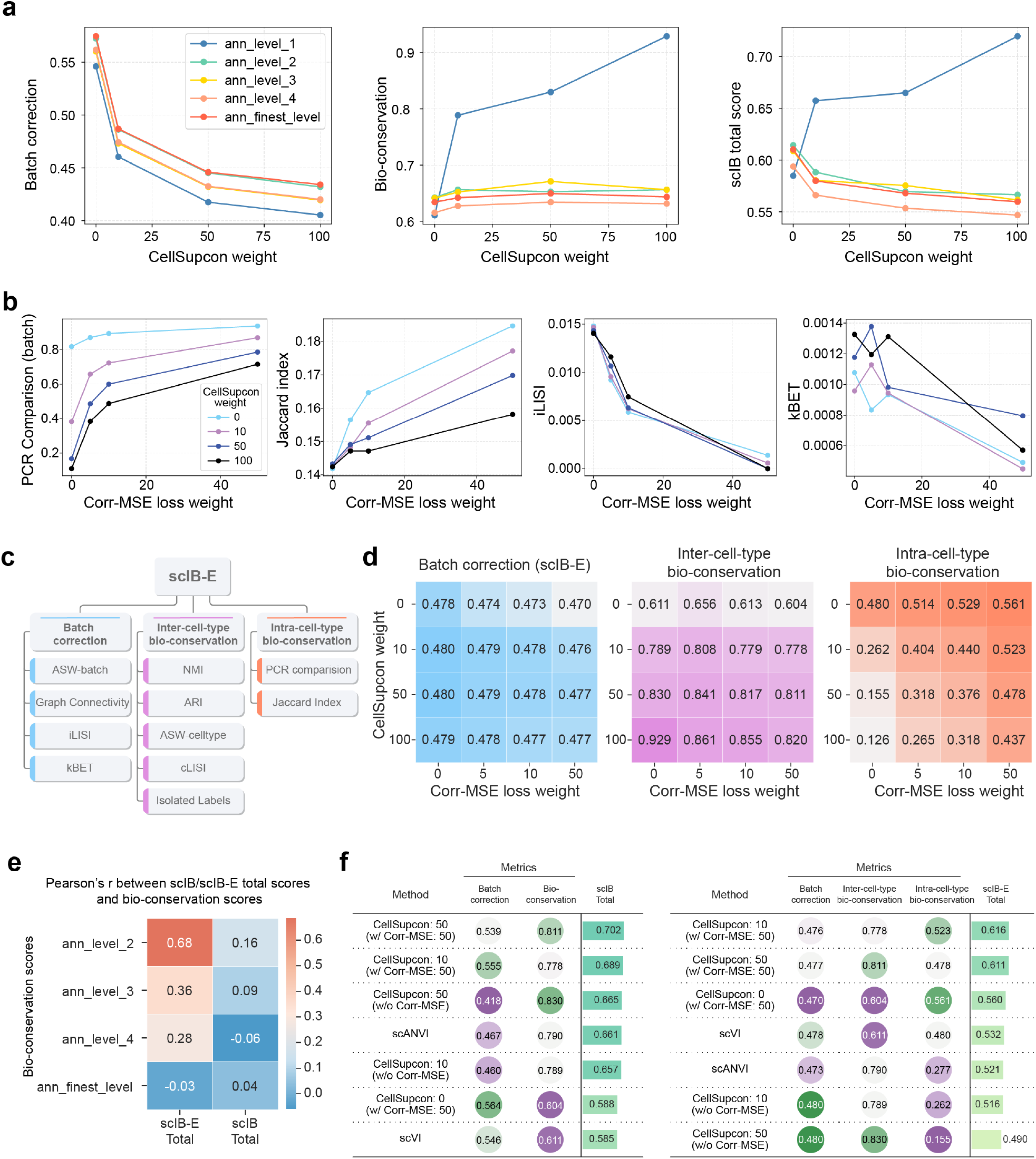
Comprehensive benchmarking of single-cell integration using extended scIB metrics. **a**, Line plots showing scIB metrics for the HLCA dataset integration results across varying CellSupcon loss weights: batch correction (left), biological conservation (middle), and total scIB score (right). The level-1 cell annotation was used for model training, while evaluations were performed with different levels of cell annotations. Colors indicate the levels of cell-type annotation used for evaluation. **b**, Line plots depicting trends in PCR comparison-batch, Jaccard index, iLISI, and kBET indices for HLCA dataset integration results with varying CellSupcon and Corr-MSE loss weights. Colors represent different CellSupcon weights. **c**, Schematic representation of the scIB-E metrics, designed for comprehensive single-cell integration benchmarking. **d**, Heatmaps showing scIB-E scores for batch correction (left), inter-cell-type biological conservation (middle), and intra-cell-type biological conservation (right) across different configurations of Corr-MSE and CellSupcon weights in the HLCA dataset. **e**, Heatmap showing Pearson correlation coefficients of scIB-E total scores (left) and scIB total scores (right) of HLCA dataset integration results using level-1 cell annotation, across paired bio-conservation scores estimated by different level cell annotations. **f**, Tables summarizing scIB (left) and scIB-E (right) metrics for HLCA dataset integration results with varying CellSupcon and Corr-MSE loss weight configurations, alongside baseline performances of scVI and scANVI. A score of 1 represents optimal performance.

To address this limitation, we introduced a correlation-based loss function, Correlation Mean Squared Error (Corr-MSE) Loss, designed to maintain the correlation similarity of single cells before and after data integration within each batch (Methods). Biological conservation scores that were evaluated at various annotation levels confirmed that this loss enhances the preservation of intra-cell-type biological variation (Supplementary Fig. 4a). Since the PCR comparison-batch index reflects the intra-cell-type biological variation, we also introduced a Jaccard index to quantify the ratio of edge connections among single cells before and after single-cell integration within a global kNN graph of each batch (Methods). We evaluated changes in the PCR comparison-batch index, Jaccard index, and other batch correlation metrics, iLISI and kBET, by varying the weights of CellSupcon and Corr-MSE losses. Empirical results demonstrated that increasing the Corr-MSE weight enhances the conservation of intra-cell-type biological conservation while restraining batch correlation metrics focused on local cell connections (Fig. 4b, Supplementary Table 7). In contrast, increasing the CellSupcon weight primarily improved the conservation of inter-cell-type information.

### Extended scIB metrics for comprehensive single-cell integration benchmarking

The scIB metrics established a set of benchmarks for single-cell integration tasks, focusing on batch correction and biological conservation. While these metrics are effective for method benchmarking, they have limitations in assessing cell-label-free intra-cell-type biological conservation. To address this, we developed an extended version, scIB-E, which encompasses three categories: batch correction, inter-cell-type biological conservation, and intra-cell-type biological conservation (Fig. 4c, Methods). Specifically, PCR comparison-batch and Jaccard indices were introduced to evaluate intra-cell-type conservation. To assess the effectiveness of the new metrics across different methods, we compared scores across these categories using various Corr-MSE and CellSupcon weight configurations applied to the single-cell lung atlas dataset (Fig. 4d). As previously noted, Corr-MSE primarily enhances intra-cell-type biological information conservation, while CellSupcon predominantly influences inter-cell-type bio-conservation (Supplementary Table 7). Additionally, we analyzed the total scores of scIB and scIB-E metrics to estimate multi-layer cell biological conservation estimation, the scIB-E total score exhibited higher Pearson correlation coefficients, indicating its superior ability to capture comprehensive biological variation in single-cell data (Fig. 4e, Supplementary Fig. 4b). Furthermore, compared to the original scIB metrics, scIB-E provided more stable batch correction scores and a clearer distinction between inter- and intra-cell-type bio-conservation scores (Fig. 4f). These findings suggest that a balanced combination of Corr-MSE and CellSupcon weights optimizes single-cell integration analysis, highlighting the advantages of scIB-E in advancing biological conservation assessment.

We then employed the scIB-E metrics to re-evaluate our previous benchmarking analysis involving different loss functions (Fig.2) and to assess the effects of Corr-MSE loss on single-cell integration. The updated scIB-E metrics revealed that the Domain class triplet loss consistently outperformed other designs, both with and without Corr-MSE regulation, while the Level-2 IRM loss demonstrated substantial performance advantages under less stringent regulation (Fig. 5a, Supplementary Figs. 5-7). By comparing index changes across all methods with and without Corr-MSE regulation, we confirmed that this regulation enhances intra-cell-type biological conservation by 6.61 ± 5.49 %, while slightly reducing the batch correction index by -4.02 ± 1.84 % and having no significant effect on inter-cell-type biological conservation (Fig. 5b). Overall, Corr-MSE regulation improves the scIB-E total index by 1.43 ± 1.71%, suggesting that this approach enhances single-cell integration by better preserving comprehensive biological variation.

**Fig 5.**
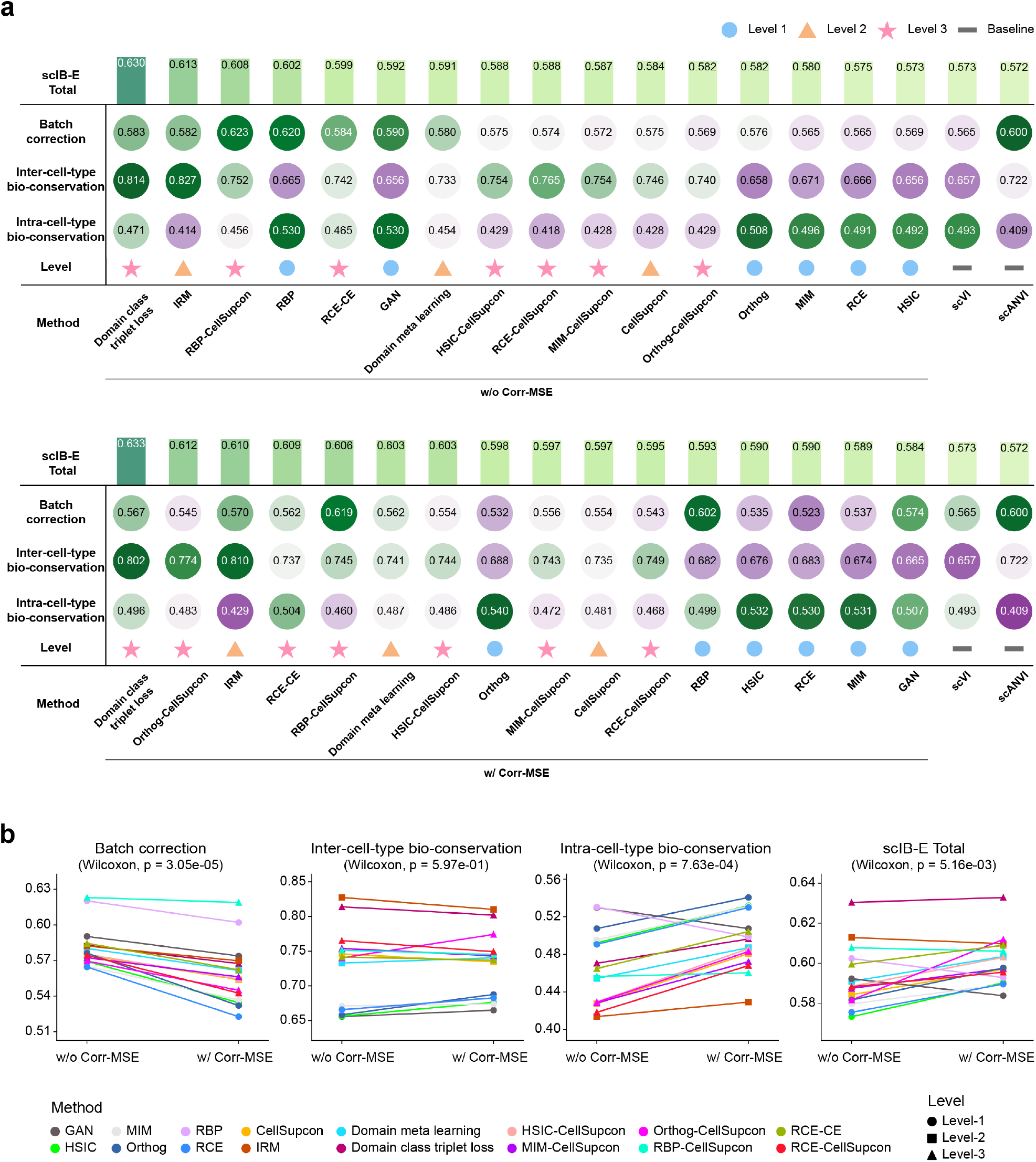
Assessing multi-level loss functions for single-cell integration using scIB-E metrics. **a**, Tables summarizing the average scIB-E scores across three datasets for multi-level loss functions, as described in **Fig. 2**. Results are shown without Corr-MSE loss (top) and with Corr-MSE loss (bottom). Methods are ranked in descending order by total score, with a score of 1 indicating optimal performance. **b**, Paired line plots illustrating average scIB-E metrics across three datasets for batch correction, inter-cell-type biological conservation, intra-cell-type biological conservation, and total score, evaluated with and without Corr-MSE loss across multi-level loss functions. Colors represent distinct methods, and shapes denote different levels. Statistical significance was assessed using a paired Wilcoxon test, with p-values labeled.

### Enhanced single-cell integration methods for preserving biological information

To further evaluate single-cell integration methods by optimizing on preserving biological variation, we analyzed a multi-layer annotated single-cell dataset from the Human Fetal Lung Cell Atlas^29^. Our analysis compared scVI, scANVI, and methods incorporating Domain class triplet loss with or without Corr-MSE loss for their integration performance. The Domain class triplet loss was identified as the top-performing method for single-cell integration from our previous analysis, while the Corr-MSE loss was applied to assess its impact on intra-cell-type variation regulation. For model training, broad cell type annotations were utilized as known cell labels, and UMAP projections were employed to visualize the developmental lung cell representations learned by different methods (Fig.6a, Supplementary Fig. 8a). Performance was assessed using scIB-E metrics across three sub-categories. The Domain Class Triplet Loss outperformed baseline scANVI in preserving both inter and intra-cell-type biological structures. In contrast, scVI, which lacks cell label constraints, achieved the highest intra-cell-type bio-conservation score. Notably, integrating Corr-MSE Loss further enhanced intra-cell-type biological variation preservation, resulting in optimized single-cell integration performance (Fig. 6b).

**Fig 6.**
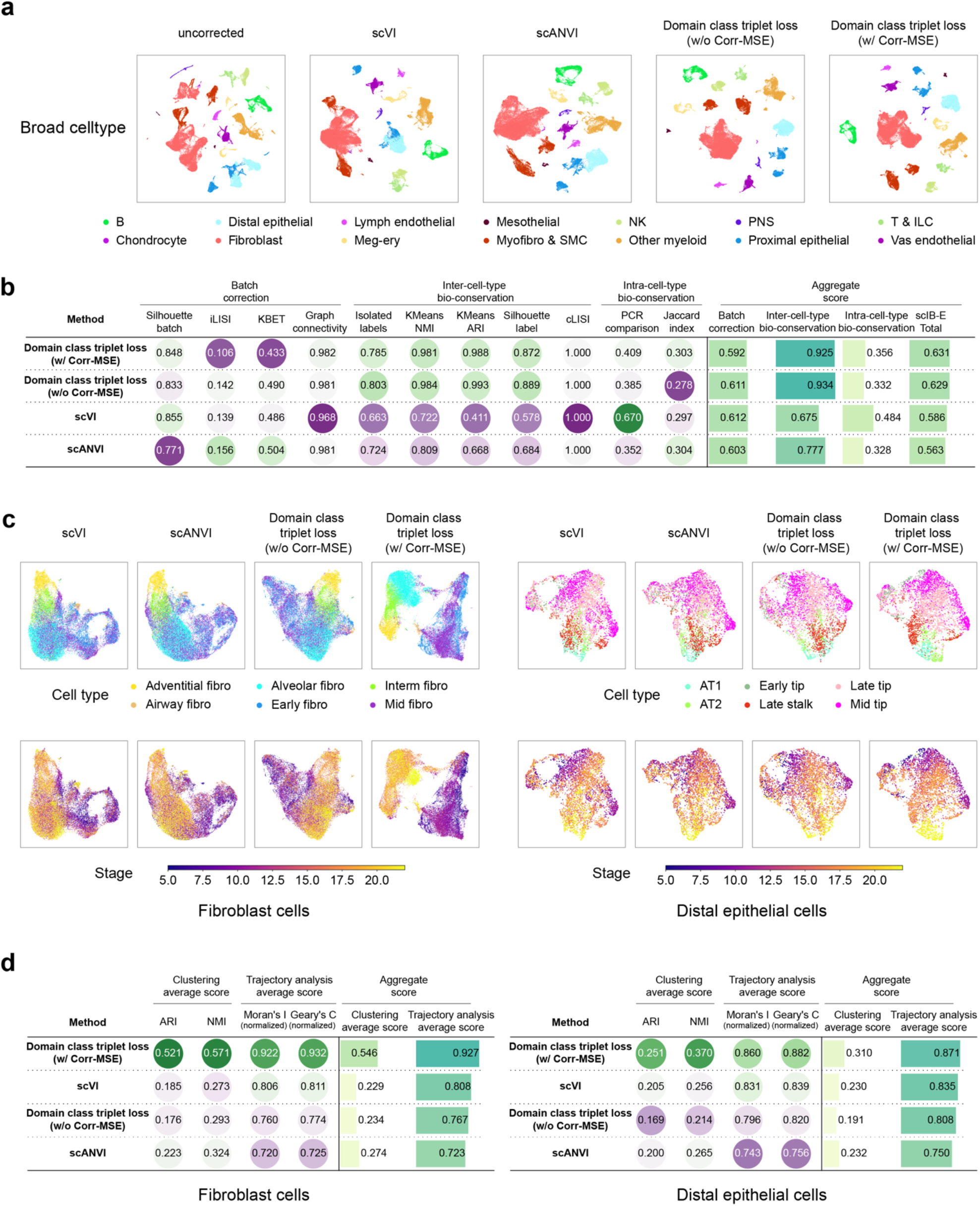
Enhanced methods for biological preserving single-cell integration. **a**, UMAP visualization of broad cell-type annotations for the Human Fetal Lung Cell Atlas, comparing the integration results of scVI, scANVI, and Domain class triplet loss methods with and without Corr-MSE loss. **b**, Table summarizing the scIB-E metrics for integration results as shown in (a). **c**, UMAP visualization of learned single-cell embeddings for fibroblast (left) and distal epithelial cells (right) from the Human Fetal Lung Cell Atlas across different integration methods as in (a), cells are colored by high-resolution cell labels (top) and developmental stages (bottom). **d**, Tables summarizing clustering and developmental trajectory conservation performance of learned single-cell embeddings for fibroblast (left) and distal epithelial cells (right) as in (c). Methods are ranked in descending order based on aggregated scores.

For a detailed examination of biological variation preservation, we subsetted learned embeddings of fibroblast and distal epithelial cells across methods (Fig. 6a). To assess the biological variation within these embeddings, we focused on high-resolution cell labels and developmental stages of selected major cell types (Fig. 6c, Supplementary Fig. 8b). Clustering performance was evaluated using ARI and NMI indices, while developmental trajectory conservation was assessed with Moran’s I and Geary’s C indices (Supplementary Table 10, Methods). Methods like scANVI and Domain Class Triplet Loss without Corr-MSE Loss relied on broad cell labels as biological constraints, limiting their ability to conserve intra-cell-type variation (Fig. 6d). In contrast, scVI achieved higher scores in developmental trajectory conservation (Fig. 6d). Furthermore, the integration of Domain Class Triplet Loss with Corr-MSE Loss demonstrated superior performance in local cell representation learning (Fig. 6d), as reflected in its high scores for broad cell type-level biological conservation (Fig. 6b). These findings highlight the versatility of the deep learning framework for single-cell integration. By leveraging different loss designs, this framework effectively balances comprehensive biological variation conservation and the removal of unwanted signals, achieving optimal performance for single-cell integration tasks.

## Discussion

In this study, we presented a multi-level benchmarking framework for single-cell data integration, leveraging deep learning to evaluate diverse loss function designs and information regularization strategies. The benchmarking results illustrate the potential of targeted loss functions to effectively balance batch effect removal and biological conservation, thereby improving single-cell data integration quality. Notably, we developed an extended set of metrics, scIB-E, that comprehensively captures both inter and intra-cell-type biological conservation, addressing limitations in traditional single-cell integration assessments.

Our findings indicate that the performance of integration methods is highly influenced by the level of information regularization applied. Stronger regularization, such as the CellSupcon^33^ loss, improves biological conservation at inter-cell-type levels but can compromise intra-cell-type information and lead to overcorrection^42^. This trade-off emphasizes the importance of carefully balancing these objectives. For instance, we demonstrated that combining Corr-MSE with CellSupcon^33^ could achieve more nuanced integration, preserving intricate biological variations while reducing batch effects. Additionally, employing the Domain Class Triplet loss^36^ in level-3 integration emerged as an effective strategy for robust biological conservation across varying cell-type resolutions. These findings suggest that combining different loss strategies can yield synergistic effects, providing researchers with a versatile toolkit for achieving specific integration goals in diverse biological contexts.

The extension of benchmarking metrics to assess intra-cell-type biological conservation provided valuable insights into the limitations of existing methods. For instance, the conventional scIB^27^ metrics often failed to capture subtle biological variations within annotated cell types, potentially overlooking biologically significant features such as disease-specific subtypes or micro-populations. Moreover, batch correction evaluations reliant on metrics like iLISI and kBET primarily focus on local cell neighbors, lacking the ability to preserve structural biological variation within batches. Including PCR comparison and Jaccard indices offers complementary approaches for evaluating the conservation of biological information within integrated cell embeddings. Additionally, by incorporating the Corr-MSE loss to maintain correlation similarities within each batch, our analyses demonstrated improved conservation of biological variation.

Using the multi-layer annotated lung atlases, we assessed various methods for intra-cell-type biological conservation, leveraging high-resolution single-cell labels and biological statuses not used in model training as proxies for intrinsic biological variation. The distinction between inter- and intra-cell-type bio-conservation scores offered by scIB-E demonstrated its superior capacity for comprehensively evaluating single-cell data integration. However, despite the improvements in preserving biological variation, the intra-batch biological signal can still be obscured by technical noise or other unwanted sources of variation. Uncovering the true biological signal within a single batch remains a significant challenge. This limitation highlights the need for further development of methods that can better disentangle true biological information from batch-specific artifacts, ensuring that the integration process does not inadvertently remove or obscure important biological signals within a single batch. Additionally, emerging approaches that aimed to isolate condition-specific biological signals were also introduced, such as contrasiveVI^43^ and DA-seq^44^.

In conclusion, our benchmarking framework and extended metrics offer a more comprehensive approach to evaluating single-cell data integration tasks, particularly in the context of preserving biological information. The development and evaluation of Corr-MSE and other tailored loss functions provide new tools for optimizing single-cell data integration, enhancing our ability to uncover complex biological processes. Future work could expand these frameworks to other single-cell modalities or experimental factors, enhancing the extraction of true biological insights. Our work underscores the value of flexible deep learning frameworks and well-designed loss functions in pushing the boundaries of single-cell data analysis, contributing to the creation of high-quality single-cell atlases that can facilitate deeper biological insights.

## Methods

### Baseline methods

#### scVI

Single-cell Variational Inference (scVI)^19^ is a probabilistic framework for the analysis and representation of scRNA-seq data. The method uses a zero-inflated negative binomial (ZINB) distribution to model gene expression and combines a variational autoencoder with a Bayesian hierarchical model to capture both biological and technical variations. scVI generates low-dimensional embeddings that can be applied to tasks such as clustering, differential expression analysis, and batch correction.

#### scANVI

Single-cell ANnotation using Variational Inference (scANVI)^21^ is a semi-supervised extension of scVI. scANVI incorporates labeled data into the generative model, guiding the latent space with cell-type labels for enhanced data integration.

### Loss designs

#### GAN

Generative Adversarial Network (GAN)^30^ is an adversarial framework that involves a generator and a discriminator engaged in a min-max optimization. The generator creates synthetic samples, while the discriminator distinguishes between real and synthetic ones, with the goal of improving the generator’s ability to produce realistic samples. We adopt the GAN loss design from scGAN^24^, tailored for batch effect removal in scRNA-seq data. This loss uses a generator to produce cell embeddings and a discriminator to predict batch labels, generating batch-independent embeddings through adversarial training. The optimization process is described by equation (1):

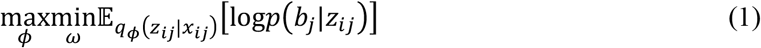

where *x*_*ij*_ represents the scRNA-seq profile of cell *i* from subject *j, b*_*j*_ is the corresponding batch label, *z*_*ij*_ is the latent embedding generated by the encoder parameterized by ϕ, and the discriminator, parameterized by ω, predicts *b*_*j*_ from *z*_*ij*_.

#### HSIC

Hilbert-Schmidt Independence Criterion (HSIC)^25^ is a non-parametric statistical test that measures the dependence between two random variables using kernel methods. HSIC computes the Hilbert-Schmidt norm of the cross-covariance operator in reproducing kernel Hilbert spaces (RKHS) to quantify the dependence between variables. We minimize the HSIC loss to ensure that cell representations are independent of batch information. The HSIC measure is defined by equation (2):

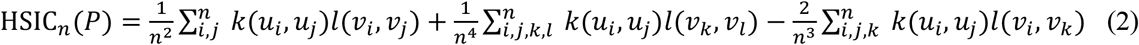

where *P* represents the joint probability distribution of the random variables, (*u*_*i*_, *v*_*i*_) are independent and identically distributed (iid) copies of the random variables *u* and *v, k* and *l* are kernel functions.

#### Orthog

Orthogonal Projection Loss (Orthog) is a statistical approach designed to reduce the correlation between two sets of embeddings by applying orthogonal constraints. This loss is minimized to enforce orthogonality between cell embeddings and batch embeddings, effectively disentangling biological signals from technical variations. The loss is formulated as the sum of the squared elements of the covariance matrix:

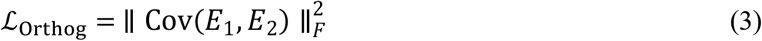

where *E*_1_ and *E*_2_ represent the two sets of embeddings, and Cov(*E*_1_, *E*_2_) denotes the covariance matrix between them.

#### MIM

Mutual Information Minimization (MIM) is an information-theoretic method that reduces the mutual information (MI) between two variables. MI quantifies the amount of information obtained about one variable through another. The definition of MI between variables *x* and *y* is in equation (4):

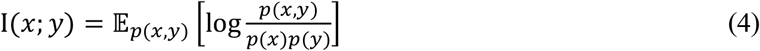

We used the Contrastive Log-ratio Upper Bound (CLUB)^45^ estimator to approximate the upper bound of MI by treating it as a divergence between joint and product distributions. To minimize the MI between cell representations and batch information, we apply the sampled vCLUB (vCLUB-S) estimator, which employs a negative sampling strategy to reduce computational complexity, It samples a single negative pair 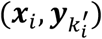 for each positive pair (***x***_*i*_, ***y***_*j*_), where 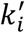 is uniformly chosen from the set {1,2, … , *N*}, excluding *i*. The MI is then estimated in equation (5):

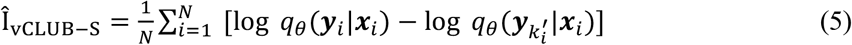

where *N* is the number of samples, and *θ* represents the parameters of the variational approximation.

#### RBP

Reverse Backpropagation (RBP)^31^ is a domain adaptation method that leverages a Gradient Reversal Layer (GRL) to learn domain-invariant representations. During forward propagation, the GRL functions as an identity transformation. During backpropagation, it reverses the gradient by multiplying it with −λ, where λ is a fixed meta-parameter:

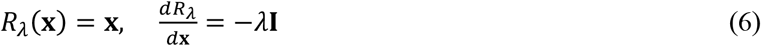

where **I** is the identity matrix. This loss helps eliminate batch-specific signals from the learned representations.

#### RCE

Reverse Cross-Entropy (RCE)^32^ encourages a uniform probability distribution across incorrect classes, introducing ambiguity to enhance the model’s robustness against label noise. We use this loss to distribute labels evenly across batches, reducing batch-specific variations. The RCE loss is defined:

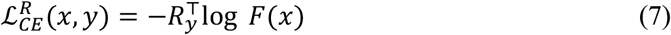

where *x* is the input feature, *y* is the target label. *F*(*x*) represents the model output as a probability value or confidence score. *R*_*y*_ is the reverse label vector, where the *y*-th element is zero, and all others are 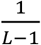, with *L* as the number of labels.

#### CellSupcon

Cell Supervised contrastive learning (CellSupcon) applies the principles of Supervised contrastive learning^33^, leveraging label information to optimize contrastive learning. In this work, cell-type labels are used as class labels, where samples from the same cell type are treated as positives, and those from different cell types are treated as negatives. The CellSupcon loss function incorporates multiple positives and negatives for each anchor sample, eliminating the need for hard negative mining. The loss is defined as:

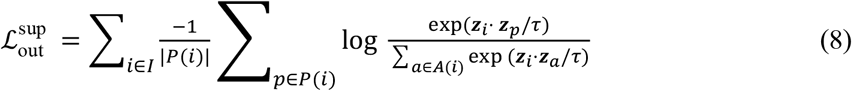

where *P*(*i*) represents the set of indices for positive samples, and *A*(*i*) comprises all other indices except *i*. ***z***_*i*_ is the feature vector of the anchor sample, ***z***_*p*_ is the feature vector of a positive sample from the same class, and ***z***_*a*_ is the feature vector of any other sample in the mini-batch. The temperature parameter τ is used for normalization.

#### IRM

Invariant Risk Minimization (IRM)^34^ is a method that enhances model generalization by learning features that remain invariant across different environments. It defines a data representation Φ: 𝒳 → ℋ that enables an optimal classifier *w*: ℋ → 𝒴 to perform consistently across all environments. The goal is to minimize the risk 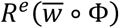 for each environment *e* to ensure stable correlations with the target variable, regardless of environmental variations. IRM is formulated as:

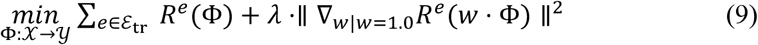

where *ε*_tr_ is the set of training environments, Φ is the entire invariant predictor, *w* = 1.0 is a fixed “dummy” classifier, and the gradient norm penalty evaluates the classifier’s optimality in each environment. The regularization parameter λ ∈ [0, ∞) balances predictive power (an empirical risk minimization term) and the invariance of the predictor 1 ⋅ Φ(*x*). We apply IRM by treating batch information as environmental variables and learning invariant features across batches to improve model generalization.

#### Domain meta-learning

Domain meta-learning^35^ is a method designed to improve model adaptability and generalization of models across different domains. It consists of two phases: meta-train and meta-test. In the meta-train phase, the model is trained on labeled data from source domains to learn lower-dimensional representations that effectively predict class labels. This is achieved using a feature extractor defined by 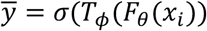, where *F* _θ_ (*x*) is the feature extractor for 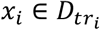, *T*_*ϕ*_ is a task-specific module, and *σ* is the softmax activation. The task-specific loss is defined in equation (10):

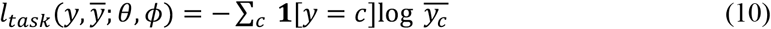

The meta-train phase loss and parameter updates are defined in equations (11 and 12):

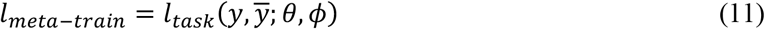

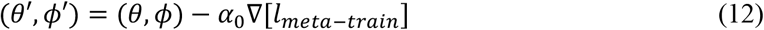

In the meta-test phase, the model learns domain-invariant representations to generalize across unseen environments. This involves aligning the geometric configuration of class centroids between domains. The centroid for class *c* in the domain *d* is defined as in equation (13):

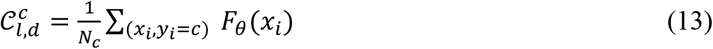

The alignment loss that measures the difference in pairwise distances between class centroids across meta-train and meta-test domains is defined in equation (14):

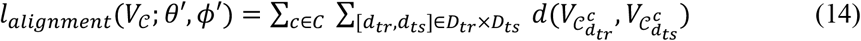

where *d*_*tr*_ ∈ *D*_*tr*_ are the meta-train domains and *d*_*ts*_ ∈ *D*_*ts*_ are the meta-test domains. The loss and the parameter update in the meta-test phase are defined in equations (15 and 16):

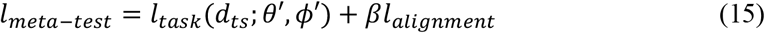

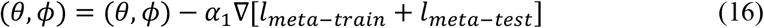

where *l*_*task*_ is derived from both labeled and unlabeled samples in *d*_*ts*_ ,and *β* is the regularization coefficient. We use domain meta learning in batch correction to improve the model’s ability to generalize across batches, ensuring consistent features are learned across different batches.

#### Domain class triplet loss

Domain class triplet loss^36^ is an extension of traditional triplet loss designed to integrate domain information for improved generalization across domains. This method modifies the triplet configuration by selecting the positive sample from the same class but a different domain, and the negative sample from the same domain but a different class. The loss function is defined in equation (17):

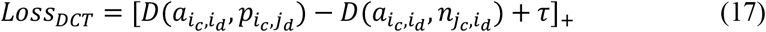

where *D* denotes the Euclidean distance, τ is a margin that specifies the separation between the positive and negative pairs. *α, p, n* represent the anchor, positive, and negative samples respectively, with *c* indicating class labels and *d* indicating domain labels. We apply this loss to integrate cell type and batch information, optimizing biological signal retention while eliminating batch effects.

#### Corr-MSE Loss

The Correlation Mean Squared Error (Corr-MSE) loss is designed to preserve global cell correlations during batch correction, minimizing the loss of cellular information. For each batch, the Pearson correlation coefficient matrices are computed for both the original PCA embeddings and the batch-corrected embeddings. The loss is computed as the mean squared error (MSE) between these two correlation matrices as described in equation (18):

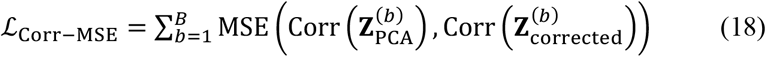

where *B* is the number of batches, 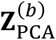 represents the original PCA embeddings for batch *b*, and 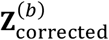 refers to the batch-corrected embeddings for batch *b* .

### Benchmark metrics

#### ASW

Average Silhouette Width (ASW)^46^ is a metric used to evaluate clustering quality by comparing within-cluster distances to between-cluster distances for each cell. ASW ranges from -1 to 1, with higher values indicating well-separated clusters, and lower values indicating poor separation or misclassification. The silhouette width *s*(*i*) for a cell *i* is calculated in eqaution (19):

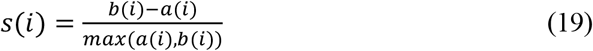

where *a*(*i*) is the mean distance to other cells in the same cluster, and *b*(*i*) is the mean distance to the nearest different cluster.

ASW assesses both cell type distinction and batch effect correction. Cell type ASW is scaled between 0 and 1 in equation (20), where higher values indicate better preservation of cell type identity after integration:

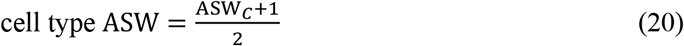

where *C* represents all cell identity labels.

Batch mixing ASW uses the absolute silhouette width *s*_batch_ (*i*) = |*s*(*i*)|, where *s*_batch_ (*i*) = 0 indicates perfect batch mixing. Batch ASW is scaled between 0 and 1, where higher values indicate effective batch integration. The batch ASW for each cell type *j* is computed in equation (21):

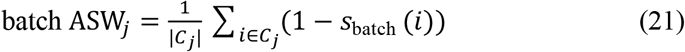

where *C*_*j*_ is the set of cells with label *j*. The overall batch ASW is the average across all cell types in equatio (22):

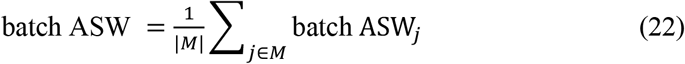

where *M* is the set of unique cell labels.

#### Graph connectivity

Graph Connectivity (GC) is a batch correction metric that measures the connectivity within label-specific subgraphs in a k-nearest neighbors (kNN) graph. The GC score ranges from 0 to 1, with higher values indicating better batch correction and well-connected cells of the same label, while lower scores suggest poor correction and fragmented subgraphs. The GC score is calculated in equation (23):

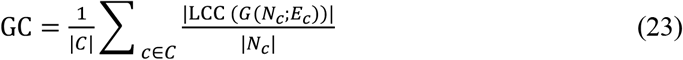

where *C* denotes the set of cell identity labels, |LCC (*G*(*N*_*c*_; *E*_*c*_))| is the number of nodes in the largest connected component for each label *c* , and |*N*_*c*_| represents the total number of nodes with that label.

#### LISI

LISI (Local Inverse Simpson’s Index)^19^ evaluates data integration by measuring diversity within kNN graphs. It has two variants: iLISI for assessing batch mixing and cLISI for evaluating cell type separation. Both scores are rescaled to 0-1, with higher values indicating better batch mixing or cell-type separation. For each node in the integrated kNN graph, iLISI is defined in equation (24):

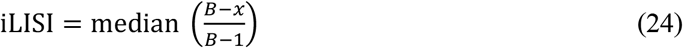

where *x* is the observed diversity in batch labels, *B* is the number of batches. cLISI is defined as in equation (25):

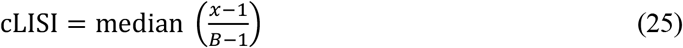

where *x* is the diversity in cell type labels, and *B* is the total number of distinct cell types.

#### kBET

kBET (k-nearest neighbor Batch Effect Test)^47^ evaluates batch effect correction by comparing local and global batch compositions within a cell’s k-nearest neighbors. The kBET score ranges from 0 to 1, where a higher score means the local batch composition closely matches the global batch composition, indicating better batch effect correction. The kBET score is calculated in equation (26):

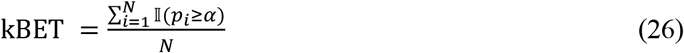

where *N* is the number of neighborhoods, 𝕀(*p*_*i*_ ≥ *α*)is the indicator function for the *i*-th neighborhood subset, equal to 1 if the p-value *p*_*i*_ is greater than or equal to the significance level *α* and 0 otherwise.

#### NMI

Normalized Mutual Information (NMI) is a statistical metric used to evaluate the similarity between two clustering results. It quantifies the extent of shared information between the two clusterings by normalizing the mutual information (MI) against the geometric mean of their entropies. The NMI score ranges from 0 to 1, where a higher value indicates better agreement between the clusterings, reflecting more accurate correspondence in assignments. The NMI is calculated in equation (27):

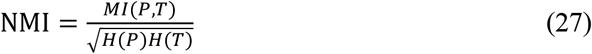

where *P* and *T* are the two sets of clustering results, *MI*(*P, T*) is the mutual information between them, and *H*(*P*) and *H*(*T*) represent the entropies, measuring the randomness within each set of clusters.

#### ARI

Adjusted Rand Index (ARI) is a metric that measures the similarity between two clustering results. It considers both correct and incorrect assignments, adjusting for chance agreement. The ARI score ranges from 0 to 1, with higher values indicating greater agreement with the ground truth. ARI is calculated in equation (28):

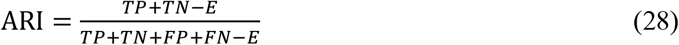

where *TP* and *TN* are the true positive and true negative pairs, *FP* and *FN* are the false positive and false negative pairs, and *E* is the expected number of random agreements calculated in equation (29):

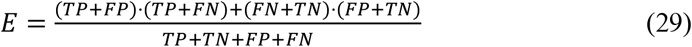

#### Isolated label score

Isolated label score is a metric designed to evaluate the effectiveness of data integration in managing cell identity labels that are present in only a few batches. It uses the average silhouette width (ASW) to assess the separation of isolated labels. The score ranges from 0 to 1, with higher values indicating better separation. The isolated label score is defined in equation (30):

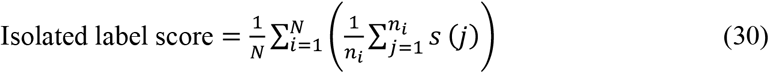

where *N* is the number of isolated labels, *n*_*i*_ is the number of samples in the *i*-th isolated label, and *s*(*j*) is the silhouette score for each sample *j*.

#### PCR comparison

Principal component regression (PCR) comparison^47^ measures the difference in explained variance before and after data integration. The total variance explained by the variable is calculated by summing the variance contributions from its influence across all principal components defined in equation (31):

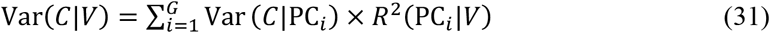

where *G* is the total number of principal components, Var(*C*|*V*) represents the variance explained by the *i*-th principal component, and *R*^2^(PC_*i*_|*V*) is the squared correlation coefficient indicating how much of the variance of the *i*-th component is explained by the variable *V*.

PCR comparison includes two metrics: PCR comparison-batch for batch removal evaluation and PCR comparison-cell for intrintic cell information evaluation. Both metrics are scaled to a range of 0 to 1, with higher values indicating better performance. The PCR comparison-batch removal is calculated in equation (32):

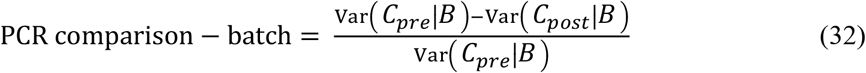

where *B* is the batch variable, V*a*r, (*C*_*pre*_ ∣ *B* ) represents the variance in the principal components explained by the batch variable before correction, and V*a*r, (*C*_*post*_ ∣ *B* ) represents the variance in the principal components after correction.

The PCR comparison-cell information retention is calculated in equation (33):

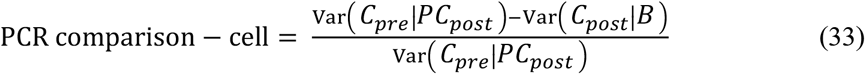

where V*a*r, *C*_*pre*_ ∣ *PC*_*post*_ 0 represents the variance in the principal components before correction, explained by the principal components after batch correction.

#### Jaccard index

Jaccard index is a global metric used to evaluate the preservation of cell information during batch correction. It measures the similarity between neighborhood structures by calculating the overlap of k-nearest neighbor graphs between the cell embeddings before and after batch correction. Jaccard index ranges from 0 to 1, with higher values indicating better preservation of neighborhood structures through the batch correction process. The Jaccard index is calculated in equation (34):

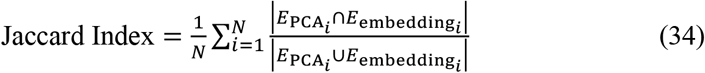

where *N* is the number of batches, 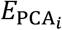 and 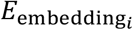 represent the sets of edges connecting the k-nearest neighbors in cell embeddings before and after batch correction, respectively.

##### Moran’s I

Moran’s I is a metric that evaluates the preservation of developmental trajectories in batch-corrected embeddings. It quantifies the global autocorrelation by measuring the correlation between developmental information and graph structures derived from batch-corrected embeddings. Moran’s I ranges from -1 to 1, with higher values indicating better trajectory preservation. The formula for Moran’s I is defined in equation (35):

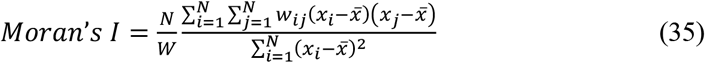

where *N* denotes the number of cells. *x*_*i*_ is the developmental information of cell *i* . 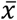 is the average developmental information. *w*_*ij*_ defines neighborhood connectivity, which is 1 if cells are neighbors and 0 otherwise. *W* is the sum of all connections. For analysis, it is normalized to [0, 1], where values closer to 1 indicate stronger preservation.

#### Geary’s C

Geary’s C evaluates local differences in developmental information among neighboring cells in batch-corrected embeddings. Its values range from 0 to 2, with lower values indicating better local similarity. The formula for Geary’s C is defined in equation (36):

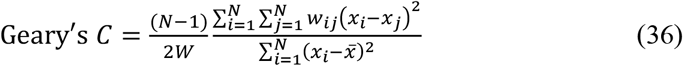

where *w*_*ij*_ is the weight between cells *i* and *j, W* = ∑_*i*,*j*_ *w*_*ij*_ is the sum of all weights. For analysis, Geary’s C is normalized to [0, 1], where higher values indicate better preservation of the developmental trajectory.

### Datasets

#### Immune

The immune dataset consists of human immune cells collected from bone marrow and peripheral blood mononuclear cells (PBMCs), sequenced using multiple platforms. It includes 33,506 cells from 10 donors and 16 cell types, collected across from five studies. The immune dataset was derived from the study^38^ and processed by selecting the 4,000 most highly variable genes.

#### Pancreas

The pancreas dataset consists of 16,382 cells collected from 9 batches, annotated and classified into 14 distinct cell types. It was obtained from the study^39^, and subset to the 4,000 most highly variable genes for further analysis.

#### BMMC

The Bone Marrow Mononuclear Cells (BMMC) dataset is created for the 2021 NeurIPS Multimodal Single-Cell Data Integration competition^40^. It contains 90,261 cells and 13,953 genes, originating from bone marrow mononuclear cells collected from multiple donors and sequenced across 12 batches^48^. The dataset includes 45 cell types and is generated using various technologies, such as RNA and protein measurements (CITE-seq) and RNA with chromatin accessibility (10x Multiome). For this analysis, the dataset focuses exclusively on transcriptomic data and is processed by selecting the 4,000 most highly variable genes.

#### HLCA

The Human Fetal Lung Cell Atlas (HLCA) is an integrated reference atlas of the human respiratory system, including lung parenchyma, respiratory airways, and the nose. We used the core dataset from the HLCA, comprising 584,944 lung cells derived from 166 samples across 107 individuals. The HLCA provides detailed annotations at multiple cell-type levels, along with comprehensive metadata. The dataset was sourced from the study^28^ and processed by selecting the 4,000 most highly variable genes.

#### Human fetal lung cell atlas

The Human fetal lung cell atlas is a dataset based on a multiomic analysis of human fetal lungs, covering 5 to 22 post-conception weeks. This dataset includes 29 batches from 12 donors and spans 8 developmental stages, encompassing 14 broad cell types and 144 newly classified cell types. Sourced from the study^29^, the dataset was processed to include only the 4,000 most highly variable genes.

### Hyperparameters and training

For model parameters, we retained the original hyperparameter settings for both the scVI and scANVI models (Supplementary Tables 1). For training, the models were trained for a maximum of 400 epochs, with a batch size of 128. Early stopping was enabled to prevent overfitting. To optimize the weight of multi-level loss function, we employed Ray Tune^37^ for hyperparameter tuning. We conducted a hyperparameter search on the immune dataset from the benchmark, selecting the set of hyperparameters that produced the best integration performance (Supplementary Tables 2). All models were implemented and trained using PyTorch (v1.13.0+cu117) with a single NVIDIA A100 GPU.

## Data availability

The immune dataset is publicly available at https://github.com/theislab/scib-reproducibility. The pancreas dataset is available at http://github.com/theislab/scPoli_reproduce. The BMMC dataset was downloaded from Gene Expression Omnibus (GEO) database unsing accession number GSE194122. The HLCA core dataset can be downloaded via cellxgene (https://cellxgene.cziscience.com/collections/6f6d381a-7701-4781-935c-db10d30de293). The Human fetal lung cell atlas was downloaded from https://fetal-lung.cellgeni.sanger.ac.uk/scRNA.html.

## Code availability

The code used for data preprocessing, data integration, and visualization in this study are available on Git-Hub at https://github.com/Chenxin-Yi/scIB-E. The code is also accessible in the Zenodo repository at https://doi.org/10.5281/zenodo.14322900.

## Acknowledgements

This study was supported by the National Key R&D Program (2023YFF1204701, 2022YFF1202101), the Self-supporting Program of Guangzhou Laboratory (SRPG22007), the CAS Research Fund (XDB38050200), Guangdong Basic and Applied Basic Research Foundation (2023B1515130008), and Department of Science and Technology of Guangdong Province (2021CX02G450). We thank technical support from the Data Science Platform of Guangzhou National Laboratory and the Bio-medical Big Data Operating System (Bio-OS).

## Author contributions

Y.L., J.L. and W.L. conceived the project. J.L. and C.Y. designed the framework and loss designs. J.C. helped data analysis. J.L. and C.Y. wrote the manuscript with contribution from all authors. Y.L. supervised the entire project. All authors read and approved the final manuscript.

## Supplementary Figs 1-8

Supplementary Fig 1: Evaluation of multi-level loss functions for single-cell integration on immune dataset.

Supplementary Fig 2: Evaluation of multi-level loss functions for single-cell integration on pancreas dataset.

Supplementary Fig 3: Evaluation of multi-level loss functions for single-cell integration on BMMC dataset.

Supplementary Fig 4: Extended scIB metrics for intra-cell-type biological conservation evaluation.

Supplementary Fig 5: Assessing multi-level loss functions for single-cell integration using scIB-E metrics on immune dataset.

Supplementary Fig 6: Assessing multi-level loss functions for single-cell integration using scIB-E metrics on pancreas dataset.

Supplementary Fig 7: Assessing multi-level loss functions for single-cell integration using scIB-E metrics on BMMC dataset.

Supplementary Fig 8: Assessing single-cell integration methods for comprehensive biological conservation.

## Supplementary Tables 1–10

Supplementary Table 1: Baseline model hyperparameters and training settings for scVI and scANVI.

Supplementary Table 2: Hyperparameters for 16 deep learning single-cell integration methods across three levels.

Supplementary Table 3: Description of the datasets used for benchmarking.

Supplementary Table 4: Detailed scIB evaluation results across three datasets for multi-level loss functions in single-cell integration.

Supplementary Table 5: Detailed scIB evaluation results for the immune dataset under varying loss regularizations.

Supplementary Table 6: Detailed evaluation of metrics for multi-level loss functions in single-cell integration: local cell-neighbors-based batch correction metrics and PCR comparison scores.

Supplementary Table 7: Detailed evaluation of metrics for multi-level loss functions, including local cell-neighbors-based batch correction metrics and PCR comparison scores.

Supplementary Table 8: Detailed scIB-E evaluation results across three datasets for multi-level loss functions in single-cell integration.

Supplementary Table 9: Detailed scIB-E metrics for enhanced methods in biologically preserving single-cell integration.

Supplementary Table 10: Downstream analysis results for enhanced methods in biologically preserving single-cell integration.

