## Supplementary Figs 1-8 for "Benchmarking deep learning methods for biologically conserved single-cell integration"

### Supplementary Figures

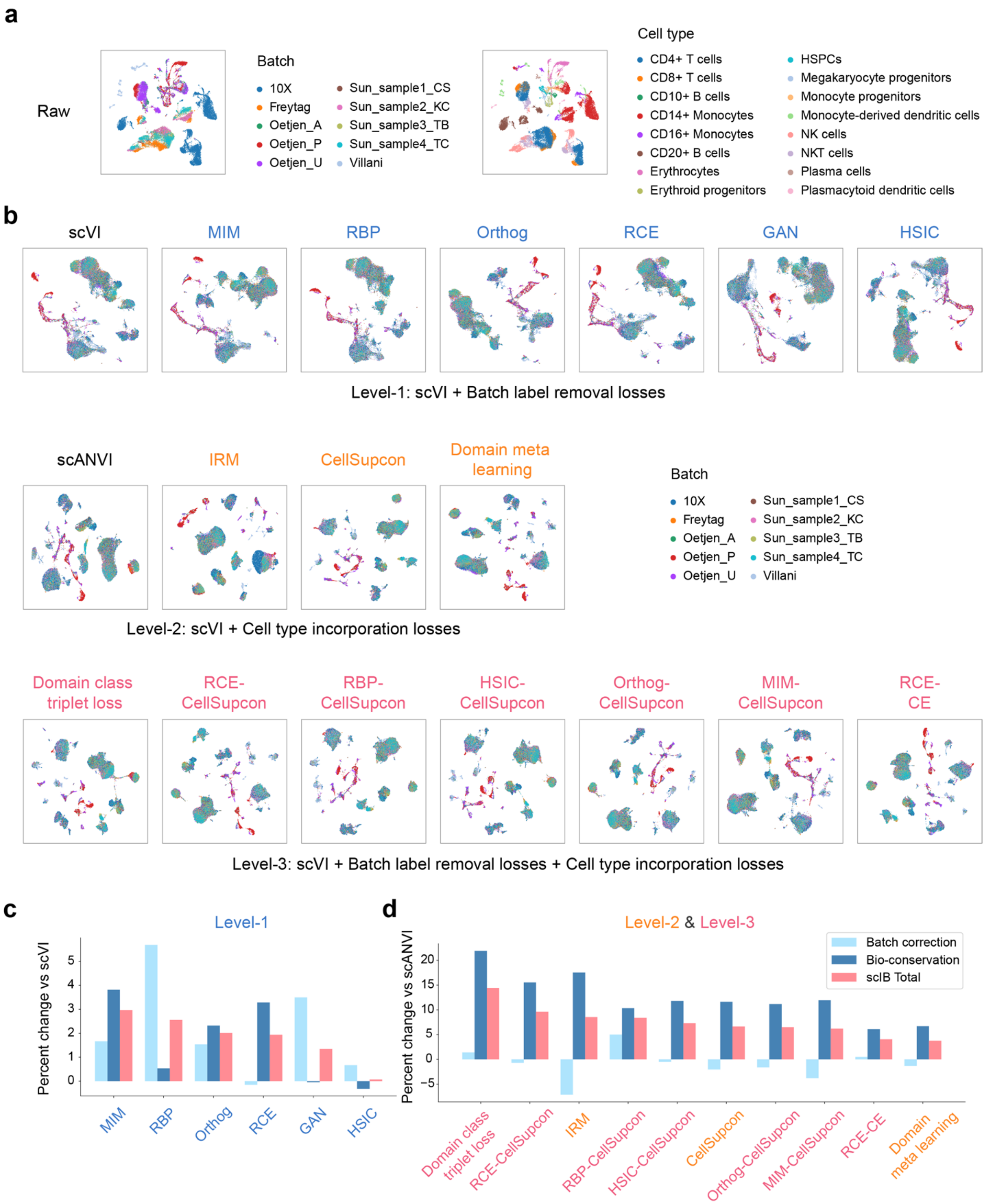

**Fig.S1 | Evaluation of multi-level loss functions for single-cell integration on immune dataset. a**, UMAP visualization of the immune dataset before integration, colored by batch label (left) and cell type (right). **b**, UMAP visualization of the integration result of immune dataset across three-level methods, colored by batch label. **c**, **d**, Comparison of scIB metrics for integration results of immune dataset across level-1 (c) and level-2/level-3 (d) methods, with scVI and scANVI as baselines, respectively. Bars represent the average percentage margins for different scores, with error bars indicating the standard error of the mean. Methods are ranked in descending order based on their total scIB scores.

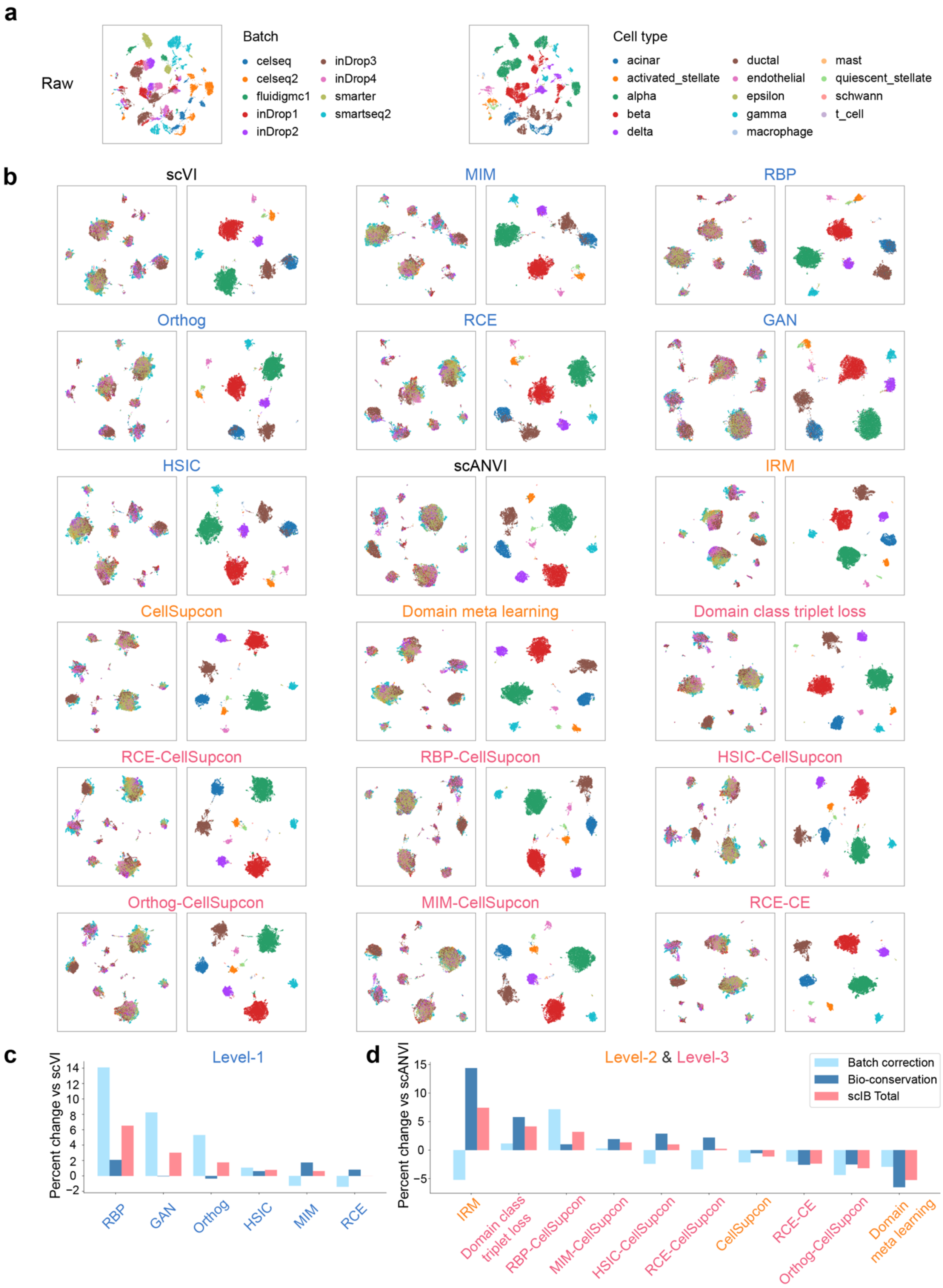
a
Raw
Batch
celseq
celseq2
fluidigmc1
inDrop1
inDrop2
inDrop3
inDrop4
smarter
smartseq2
Cell type
acinar
activated_stellate
alpha
beta
delta
ductal
endothelial
epsilon
gamma
macrophage
mast
quiescent_stellate
schwann
t_cell
b
scVI
MIM
RBP
Orthog
RCE
GAN
HSIC
scANVI
IRM
CellSupcon
Domain meta learning
Domain class triplet loss
RCE-CellSupcon
RBP-CellSupcon
HSIC-CellSupcon
Orthog-CellSupcon
MIM-CellSupcon
RCE-CE
c
Level-1
Percent change vs scVI
RBP
GAN
Orthog
HSIC
MIM
RCE
d
Level-2 & Level-3
Percent change vs scANVI
Batch correction
Bio-conservation
scIB Total
IRM
Domain class triplet loss
RBP-CellSupcon
MIM-CellSupcon
HSIC-CellSupcon
RCE-CellSupcon
CellSupcon
RCE-CE
Orthog-CellSupcon
Domain meta learning

**Fig.S2 | Evaluation of multi-level loss functions for single-cell integration on pancreas dataset. a**, UMAP visualization of the pancreas dataset before integration, colored by batch (left) and cell type (right). **b**, UMAP visualization of the integration result of pancreas dataset across three-level methods, with cells colored by batch label (left) and by cell type (right) for each method. **c**, **d**, Comparison of scIB metrics for integration results of pancreas dataset across level-1 (c) and level-2/level-3 (d) methods, with scVI and scANVI as baselines, respectively. Bars represent the average percentage margins for different scores, with error bars indicating the standard error of the mean. Methods are ranked in descending order based on their total scIB scores.

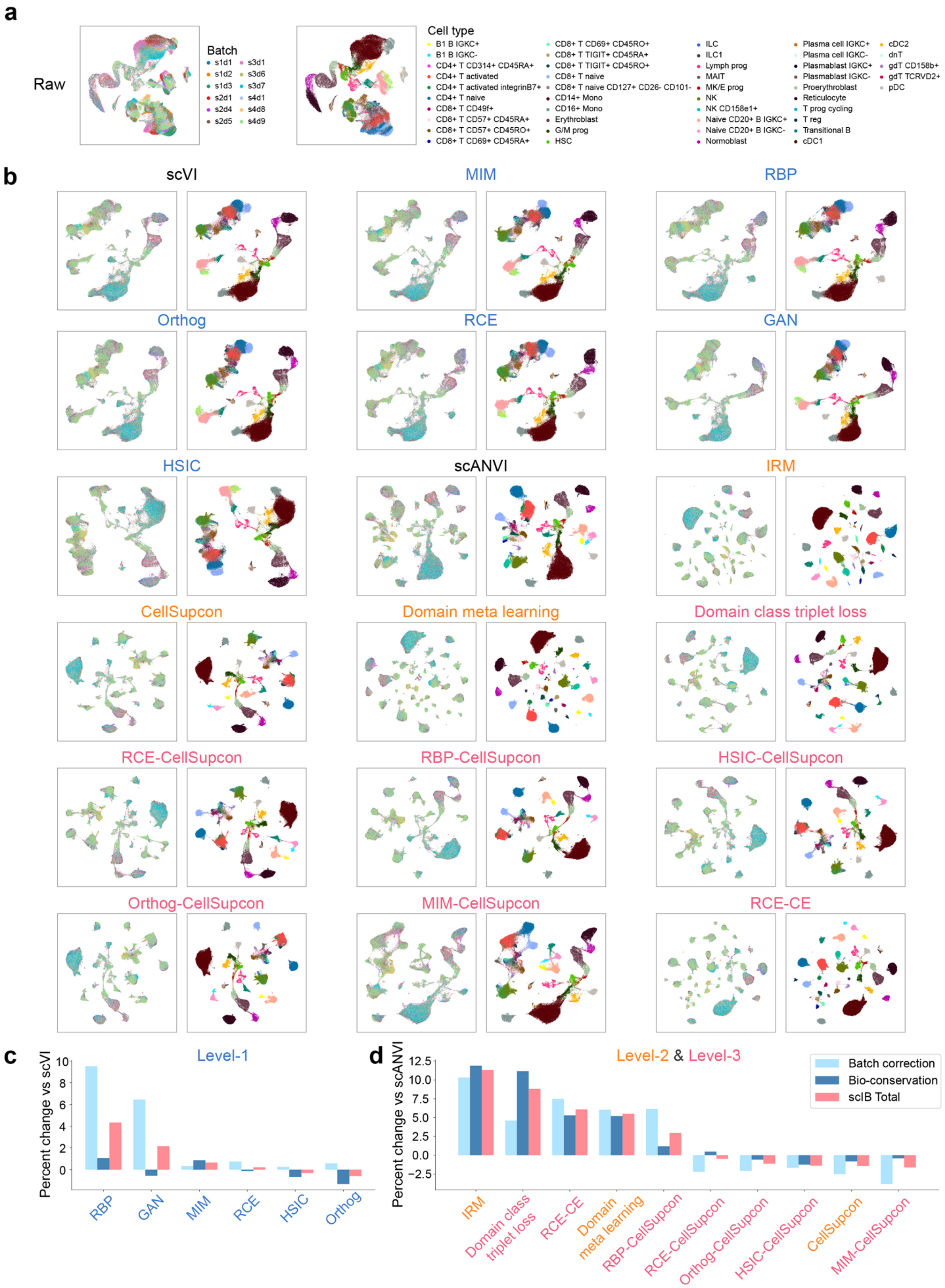

a
Raw
Batch
s1d1
s1d2
s1d3
s2d1
s2d4
s2d5
s3d1
s3d6
s3d7
s4d1
s4d8
s4d9
Cell type
B1 B IGKC+
B1 B IGKC-
CD4+ T CD314+ CD45RA+
CD4+ T activated
CD4+ T activated integrinB7+
CD4+ T naive
CD8+ T CD49f+
CD8+ T CD57+ CD45RA+
CD8+ T CD57+ CD45RO+
CD8+ T CD69+ CD45RA+
CD8+ T CD69+ CD45RO+
CD8+ T TIGIT+ CD45RA+
CD8+ T TIGIT+ CD45RO+
CD8+ T naive
CD8+ T naive CD127+ CD26- CD101-
CD14+ Mono
CD16+ Mono
Erythroblast
G/M prog
HSC
ILC
ILC1
Lymph prog
MAIT
MK/E prog
NK
NK CD158e1+
Naive CD20+ B IGKC+
Naive CD20+ B IGKC-
Normoblast
Plasma cell IGKC+
Plasma cell IGKC-
Plasmablast IGKC+
Plasmablast IGKC-
Proerythroblast
Reticulocyte
T prog cycling
T reg
Transitional B
cDC1
cDC2
dnT
gdT CD158b+
gdT TCRVD2+
pDC
b
scVI
MIM
RBP
Orthog
RCE
GAN
HSIC
scANVI
IRM
CellSupcon
Domain meta learning
Domain class triplet loss
RCE-CellSupcon
RBP-CellSupcon
HSIC-CellSupcon
Orthog-CellSupcon
MIM-CellSupcon
RCE-CE
c
Level-1
Percent change vs scVI
10
8
6
4
2
0
RBP
GAN
MIM
RCE
HSIC
Orthog
d
Level-2 & Level-3
Percent change vs scANVI
12.5
10.0
7.5
5.0
2.5
0.0
−2.5
Batch correction
Bio-conservation
scIB Total
IRM
Domain class triplet loss
RCE-CE
Domain meta learning
RBP-CellSupcon
RCE-CellSupcon
Orthog-CellSupcon
HSIC-CellSupcon
CellSupcon
MIM-CellSupcon

**Fig.S3 | Evaluation of multi-level loss functions for single-cell integration on BMMC dataset. a**, UMAP visualization of the BMMC dataset before integration, colored by batch (left) and cell type (right). **b**, UMAP visualization of the integration result of BMMC dataset across three-level methods, with cells colored by batch label (left) and by cell type (right) for each method. **c**, **d**, Comparison of scIB metrics for integration results of BMMC dataset across level-1 (c) and level-2/level-3 (d) methods, with scVI and scANVI as baselines, respectively. Bars represent the average percentage margins for different scores, with error bars indicating the standard error of the mean. Methods are ranked in descending order based on their total scIB scores.

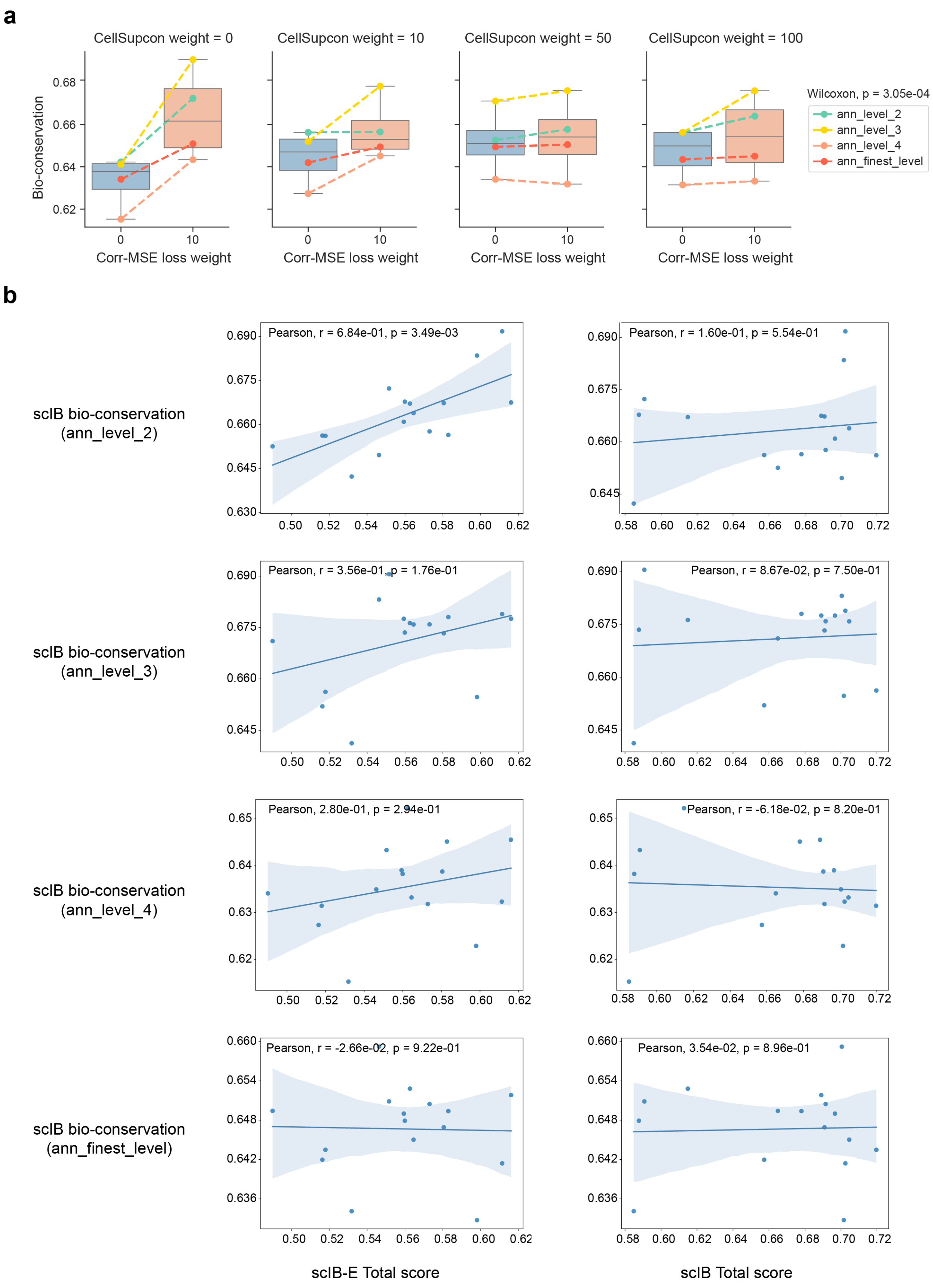
a
CellSupcon weight = 0
CellSupcon weight = 10
CellSupcon weight = 50
CellSupcon weight = 100
Bio-conservation
Corr-MSE loss weight
Wilcoxon, p = 3.05e-04
ann_level_2
ann_level_3
ann_level_4
ann_finest_level
b
scIB bio-conservation (ann_level_2)
Pearson, r = 6.84e-01, p = 3.49e-03
Pearson, r = 1.60e-01, p = 5.54e-01
scIB bio-conservation (ann_level_3)
Pearson, r = 3.56e-01, p = 1.76e-01
Pearson, r = 8.67e-02, p = 7.50e-01
scIB bio-conservation (ann_level_4)
Pearson, 2.80e-01, p = 2.94e-01
Pearson, r = -6.18e-02, p = 8.20e-01
scIB bio-conservation (ann_finest_level)
Pearson, r = -2.66e-02, p = 9.22e-01
Pearson, 3.54e-02, p = 8.96e-01
scIB-E Total score
scIB Total score

**Fig.S4 | Extended scIB metrics for intra-cell-type biological conservation evaluation. a**, Paired boxplots showing bio-conservation scores of integration results with and without Corr-MSE loss at different cell annotation levels of HLCA dataset, across varying CellSupcon loss weights. Colored lines represent different cell annotation levels. **b**, Scatter plots showing the Pearson's correlation between scIB-E total scores (left) and scIB total scores (right) of HLCA dataset integration results using level-1 cell annotation, across paired bio-conservation scores estimated by different level cell annotations, with p-values labeled. Statistical significance was assessed using a paired Wilcoxon test in (a), with p-values labeled.

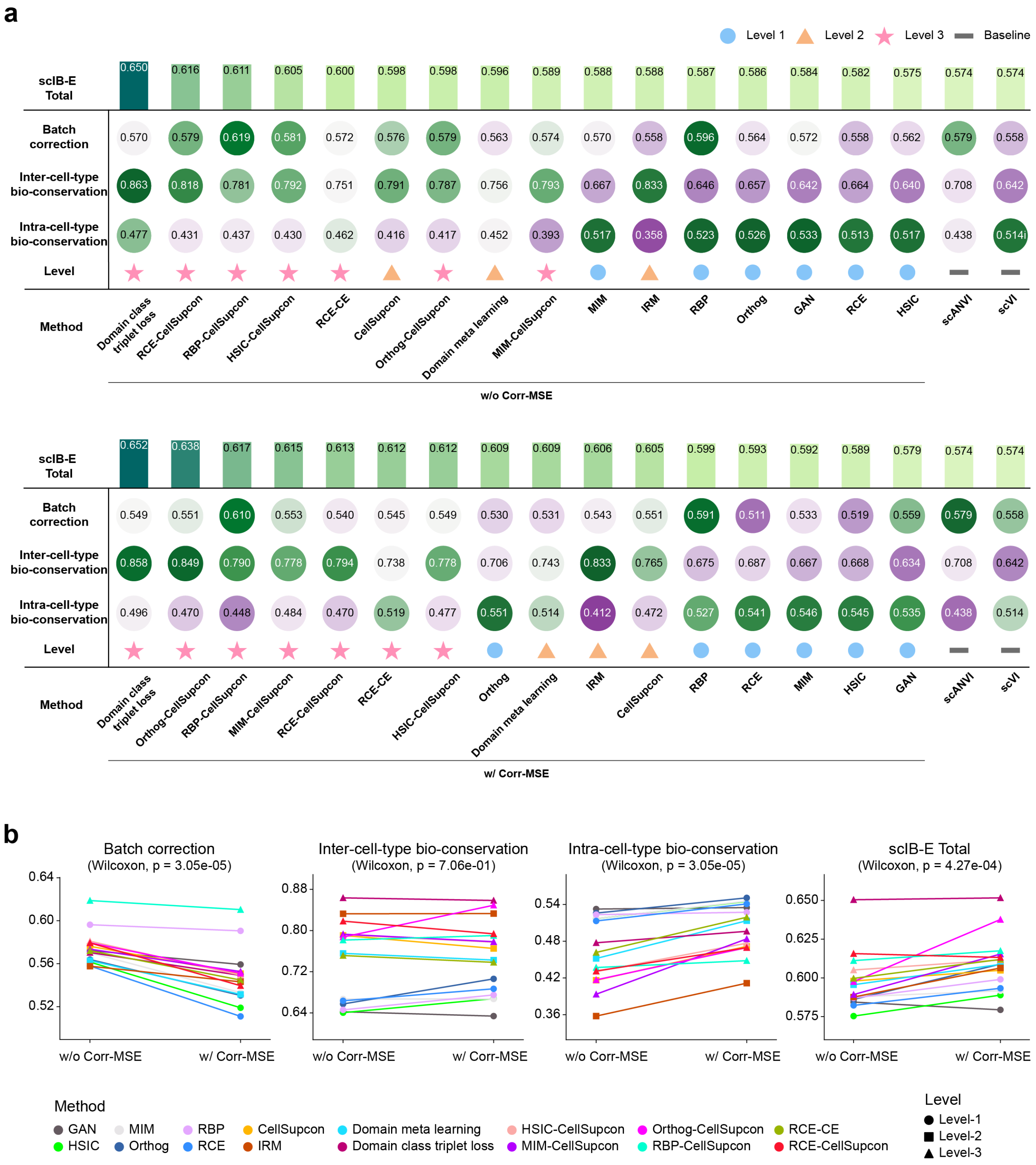


**Fig.S5 | Assessing multi-level loss functions for single-cell integration using scIB-E metrics on immune dataset. a**, Tables summarizing the scIB-E scores of the immune dataset for multi-level loss functions, as described in **Fig. 2**. Results are shown without Corr-MSE loss (top) and with Corr-MSE loss (bottom). Methods are ranked in descending order by total score, with a score of 1 indicating optimal performance. **b**, Paired line plots illustrating scIB-E metrics on immune dataset for batch correction, inter-cell-type biological conservation, intra-cell-type biological conservation, and total score, evaluated with and without Corr-MSE loss across multi-level loss functions. Colors represent distinct methods, and shapes denote different levels. Statistical significance was assessed using a paired Wilcoxon test, with p-values labeled.

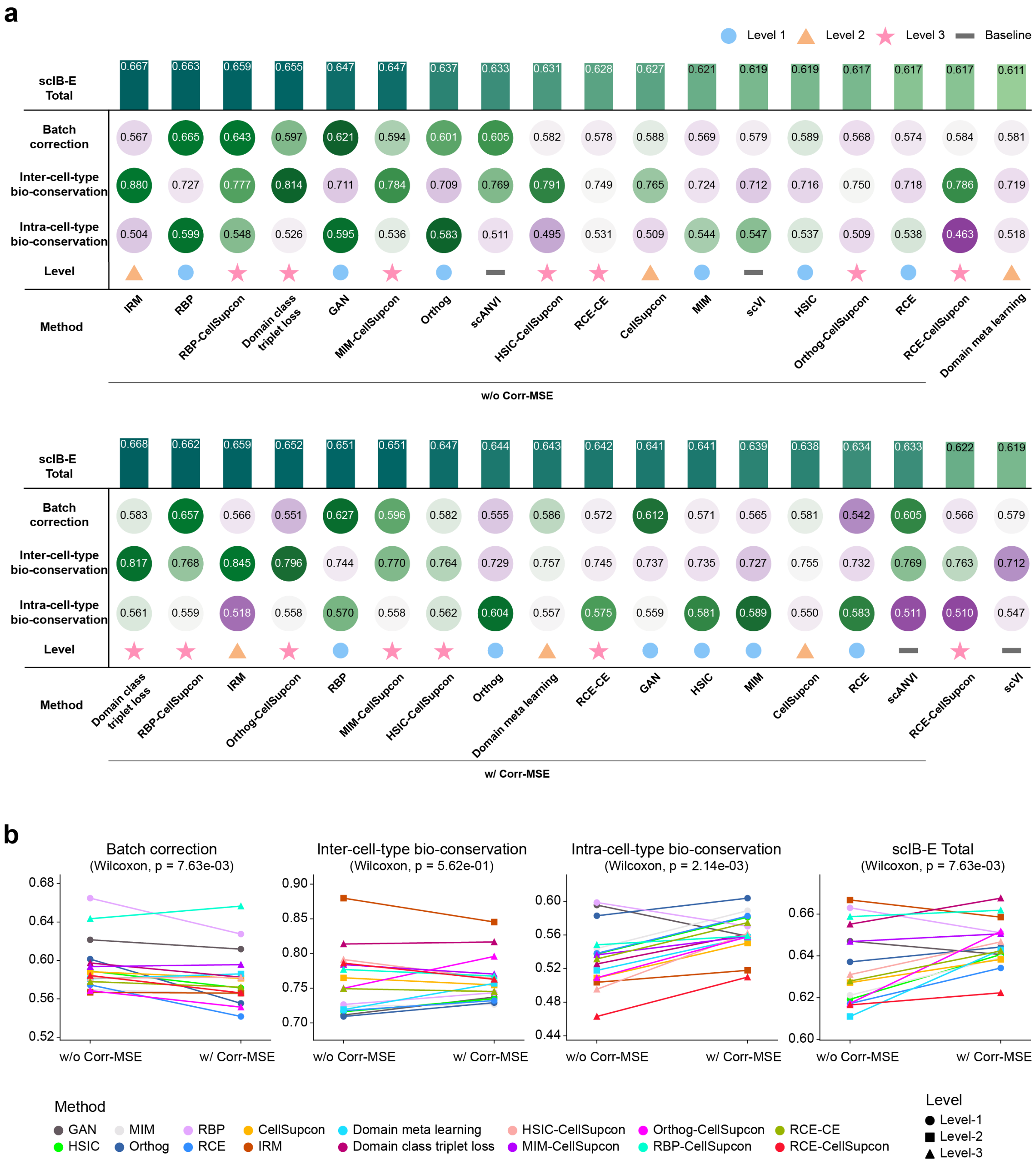


**Fig.S6 | Assessing multi-level loss functions for single-cell integration using scIB-E metrics on pancreas dataset. a**, Tables summarizing the scIB-E scores of the pancreas dataset for multi-level loss functions, as described in **Fig. 2**. Results are shown without Corr-MSE loss (top) and with Corr-MSE loss (bottom). Methods are ranked in descending order by total score, with a score of 1 indicating optimal performance. **b**, Paired line plots illustrating scIB-E metrics on pancreas dataset for batch correction, inter-cell-type biological conservation, intra-cell-type biological conservation, and total score, evaluated with and without Corr-MSE loss across multi-level loss functions. Colors represent distinct methods, and shapes denote different levels. Statistical significance was assessed using a paired Wilcoxon test, with p-values labeled.

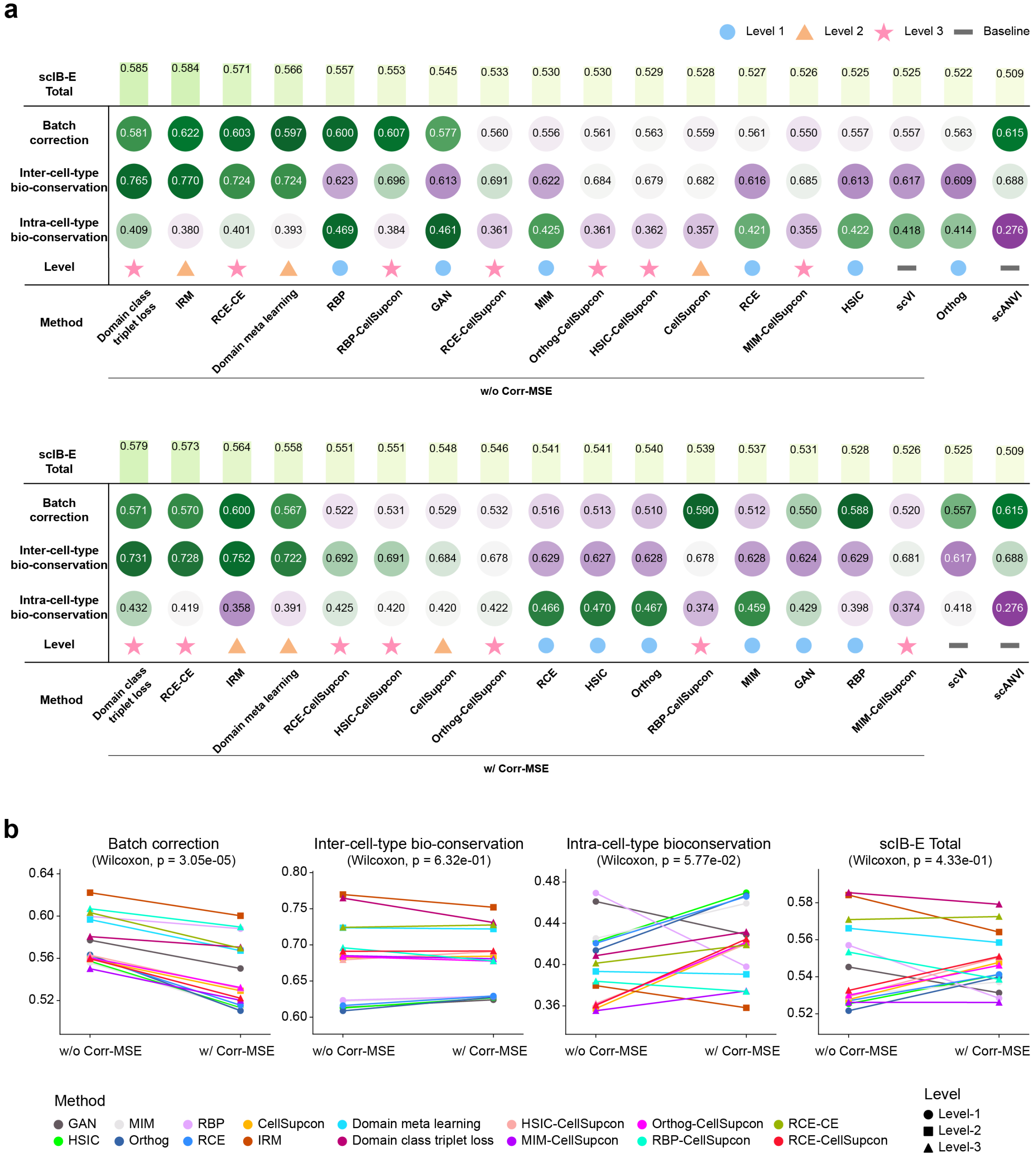


**Fig.S7 | Assessing multi-level loss functions for single-cell integration using scIB-E metrics on BMMC dataset. a**, Tables summarizing the scIB-E scores of the BMMC dataset for multi-level loss functions, as described in **Fig. 2**. Results are shown without Corr-MSE loss (top) and with Corr-MSE loss (bottom). Methods are ranked in descending order by total score, with a score of 1 indicating optimal performance. **b**, Paired line plots illustrating scIB-E metrics on BMMC dataset for batch correction, inter-cell-type biological conservation, intra-cell-type biological conservation, and total score, evaluated with and without Corr-MSE loss across multi-level loss functions. Colors represent distinct methods, and shapes denote different levels. Statistical significance was assessed using a paired Wilcoxon test, with p-values labeled.

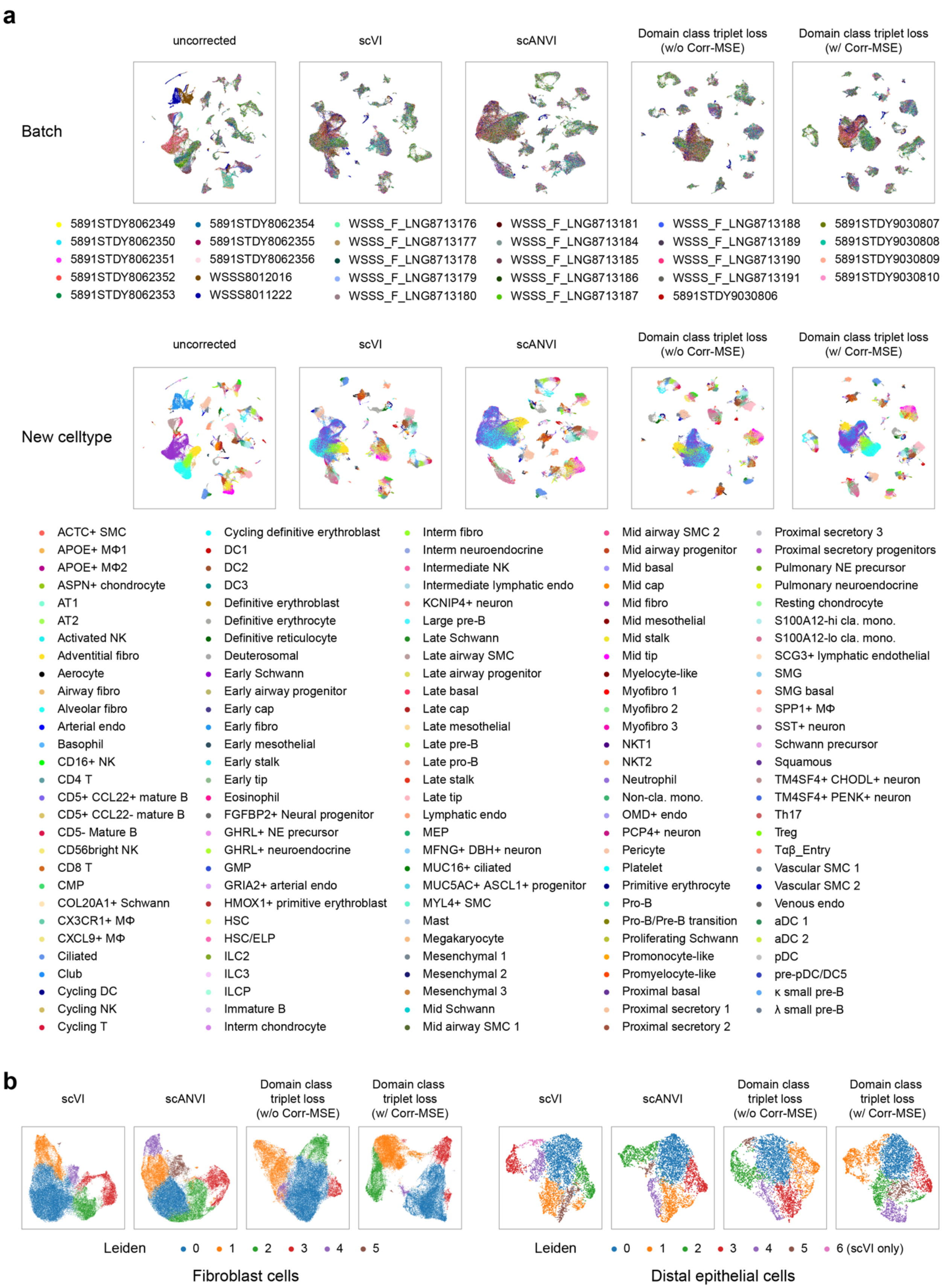
a
uncorrected
scVI
scANVI
Domain class triplet loss (w/o Corr-MSE)
Domain class triplet loss (w/ Corr-MSE)
Batch
5891STDY8062349
5891STDY8062350
5891STDY8062351
5891STDY8062352
5891STDY8062353
5891STDY8062354
5891STDY8062355
5891STDY8062356
WSSS8012016
WSSS8011222
WSSS_F_LNG8713176
WSSS_F_LNG8713177
WSSS_F_LNG8713178
WSSS_F_LNG8713179
WSSS_F_LNG8713180
WSSS_F_LNG8713181
WSSS_F_LNG8713184
WSSS_F_LNG8713185
WSSS_F_LNG8713186
WSSS_F_LNG8713187
WSSS_F_LNG8713188
WSSS_F_LNG8713189
WSSS_F_LNG8713190
WSSS_F_LNG8713191
5891STDY9030806
5891STDY9030807
5891STDY9030808
5891STDY9030809
5891STDY9030810
uncorrected
scVI
scANVI
Domain class triplet loss (w/o Corr-MSE)
Domain class triplet loss (w/ Corr-MSE)
New celltype
ACTC+ SMC
APOE+ MΦ1
APOE+ MΦ2
ASPN+ chondrocyte
AT1
AT2
Activated NK
Adventitial fibro
Aerocyte
Airway fibro
Alveolar fibro
Arterial endo
Basophil
CD16+ NK
CD4 T
CD5+ CCL22+ mature B
CD5+ CCL22- mature B
CD5- Mature B
CD56bright NK
CD8 T
CMP
COL20A1+ Schwann
CX3CR1+ MΦ
CXCL9+ MΦ
Ciliated
Club
Cycling DC
Cycling NK
Cycling T
Cycling definitive erythroblast
DC1
DC2
DC3
Definitive erythroblast
Definitive erythrocyte
Definitive reticulocyte
Deuterosomal
Early Schwann
Early airway progenitor
Early cap
Early fibro
Early mesothelial
Early stalk
Early tip
Eosinophil
FGFBP2+ Neural progenitor
GHRL+ NE precursor
GHRL+ neuroendocrine
GMP
GRIA2+ arterial endo
HMOX1+ primitive erythroblast
HSC
HSC/ELP
ILC2
ILC3
ILCP
Immature B
Interm chondrocyte
Interm fibro
Interm neuroendocrine
Intermediate NK
Intermediate lymphatic endo
KCNIP4+ neuron
Large pre-B
Late Schwann
Late airway SMC
Late airway progenitor
Late basal
Late cap
Late mesothelial
Late pre-B
Late pro-B
Late stalk
Late tip
Lymphatic endo
MEP
MFNG+ DBH+ neuron
MUC16+ ciliated
MUC5AC+ ASCL1+ progenitor
MYL4+ SMC
Mast
Megakaryocyte
Mesenchymal 1
Mesenchymal 2
Mesenchymal 3
Mid Schwann
Mid airway SMC 1
Mid airway SMC 2
Mid airway progenitor
Mid basal
Mid cap
Mid fibro
Mid mesothelial
Mid stalk
Mid tip
Myelocyte-like
Myofibro 1
Myofibro 2
Myofibro 3
NKT1
NKT2
Neutrophil
Non-cla. mono.
OMD+ endo
PCP4+ neuron
Pericyte
Platelet
Primitive erythrocyte
Pro-B
Pro-B/Pre-B transition
Proliferating Schwann
Promonocyte-like
Promyelocyte-like
Proximal basal
Proximal secretory 1
Proximal secretory 2
Proximal secretory 3
Proximal secretory progenitors
Pulmonary NE precursor
Pulmonary neuroendocrine
Resting chondrocyte
S100A12-hi cla. mono.
S100A12-lo cla. mono.
SCG3+ lymphatic endothelial
SMG
SMG basal
SPP1+ MΦ
SST+ neuron
Schwann precursor
Squamous
TM4SF4+ CHODL+ neuron
TM4SF4+ PENK+ neuron
Th17
Treg
Tαβ_Entry
Vascular SMC 1
Vascular SMC 2
Venous endo
aDC 1
aDC 2
pDC
pre-pDC/DC5
κ small pre-B
λ small pre-B
b
scVI
scANVI
Domain class triplet loss (w/o Corr-MSE)
Domain class triplet loss (w/ Corr-MSE)
Leiden 0 1 2 3 4 5
Fibroblast cells
scVI
scANVI
Domain class triplet loss (w/o Corr-MSE)
Domain class triplet loss (w/ Corr-MSE)
Leiden 0 1 2 3 4 5 6 (scVI only)
Distal epithelial cells

**Fig.S8 | Assessing single-cell integration methods for comprehensive biological conservation. a**, UMAP visualization of the Human Fetal Lung Cell Atlas as in **Fig. 6a**, cells are colored by fine-grained cell-type annotations. **b**, UMAP visualization of fibroblast (left) and distal epithelial cells (right) as in **Fig. 6c**, colored by Leiden clustering labels.
